# X-Gutor: a high-fidelity *in vitro* platform for long-term gut microbiome recovery

**DOI:** 10.64898/2026.06.09.731108

**Authors:** Qihao Li, Siyan Zhao, Jiangjian Shi, Zhiwei Liang, Yixuan Wang, Xiangpeng Wang, Jun Hu, Xiaojun Zhang, Xianyong Ma, Jing You, Guojun Shi, Zhili He, Lei Wang, Shanquan Wang

## Abstract

*In vitro* gut microbiome-culturing models are essential for studying host-microbe interactions, yet achieving high microbial recovery and long-term stability remains a long-standing challenge. Here we devise *X*-Gutor, an ecologically designed *in vitro* platform that recapitulates the monogastric digestive system with precise control of pH, redox potential, and a mucin-based biofilm matrix. Using an integrated assessment framework, microbial source tracking, a curated gut metabolism database (GutDB), and short-chain fatty acid profiling, we benchmarked gut community recovery. Building on this platform, HanGutor and PigGutor were developed to recover 96.27±4.92% and 90.48±5.95% of human and swine gut microbiota, respectively. Ecological fine-tuning revealed that redox potential drives the *Prevotella*–*Bacteroides* trade-off and that spatial biofilm structure suppresses cheater (*Succinivibrio*) growth. The *X*-Gutor provides a scalable, modular platform for gut microbiome engineering, dietary intervention, and microbiota-targeted therapy development.

## Main

The human gut microbiota acts as a metabolically active organ that integrates immunity and host metabolism. It supports nutrient fermentation, neuroimmune signaling, drug metabolism, and the production of diverse bioactive metabolites ^1–3^. Disruptions to this ecosystem are implicated in a wide range of gastrointestinal, metabolic, and systemic disorders^4^. While murine models have been the primary tool for interrogating host-microbe interactions^2,5,6^, their translational utility is constrained by fundamental differences in gut physiology and microbial composition, with species-level overlap often below 20%. These limitations, coupled with ethical concerns and throughput constraints, underscore the urgent need for robust, high-fidelity *in vitro* models capable of recapitulating the human gut microbiome^7,8^.

Current *in vitro* platforms are largely divided into bioreactors and microengineered systems^9^ (Table 1). While bioreactor platform (e.g., SHIME, TIM-2, and PolyFermS) support long-term microbial dynamics, and gut-on-chip/plate systems exploit microengineering and fluid dynamics to regulate physicochemical microenvironments and facilitate host-microbe co-culture^10,11^, both face a persistent trade-off between biological complexity and technical feasibility. Furthermore, a central limitation across existing models is the lack of a standardized framework to quantify microbiota recovery, ecological fidelity, and long-term stability ^12–14^. Current models often fail to preserve the ecological processes, such as metabolic cross-feeding and competition-cooperation trade-offs, that govern community resilience. A meta-analysis of existing literature (Fig. S1) indicates that typical microbial recovery relative to native host communities rarely exceeds 60%, with stable maintenance often limited to short time scales^10,15,16^ ^17^. Thus, there is a critical need for an ecology-informed design that moves beyond mere physicochemical tuning to actively engineer microbial interaction niches.

**Table 1.** Summary of *in vitro* culturing platforms for gut microbiomes.

| Platform | Model Type | Host Cell Co-culture | Peristaltic Movements | Biofilm/Mucus Compartment | Controllable Parameters | Stability Duration | Quantitative Fidelity Evaluation | Reference |
| --- | --- | --- | --- | --- | --- | --- | --- | --- |
| MiPro | Gut-on-plate | No | No | No | Anaerobic atmosphere, shaking | ~3 days | No | 18 |
| HuMix | Gut-on-chip | Yes | No | No | Flow, oxygen gradients | Days | No | 19 |
| Emulate Chip | Gut-on-chip | Yes | Yes | Partial | Flow, shear stress, O<sub>2</sub> gradients | Days | No | 20 |
| SHIME | Bioreactor | Yes | No | Yes | pH, retention time | Weeks | No | 21,22 |
| TIM-2 | Bioreactor | No | Yes | Yes | pH | Days–Weeks | No | 23 |
| PolyFermS | Bioreactor | No | No | Yes | pH | Weeks | No | 24 |
| MBRA | Bioreactor | No | No | No | pH, flow, anaerobic | Days | No | 25 |
| X-Gutor | Bioreactor | Optional/future compatible | Yes | Yes | pH, redox, anaerobic, biofilm structure | >180 days | Yes (>90% recovery) (community composition recovery with microbial source tracking analysis; GutDB-based functional gene recovery) | This study |

Here, we present X-Gutor, an ecologically informed platform explicitly engineered to preserve gut microbiota fidelity across community structure, functional gene capacity, and metabolic output. X-Gutor features a three-compartment reactor (stomach, small intestine, colon) with precise control of pH, redox potential, and a mucin-coated 3D-printed biofilm matrix. It employs a modular core architecture that allows for the creation of host-specific derivatives, including HanGutor (human) and PigGutor (swine), by leveraging conserved physiological features of monogastric digestive systems. To address the historic lack of reproducibility, we integrated a comprehensive, data-driven benchmarking framework (Fig. 1) comprising: (i) quantitative microbial source tracking for structural fidelity; (ii) a curated gut metabolism database (GutDB) for benchmarking functional gene capacity; and (iii) high-resolution short-chain fatty acid (SCFA) profiling to validate metabolic outputs against host-derived references. By explicitly modulating microbial competition and cross-feeding constraints, X-Gutor achieves unprecedented performance, reaching 96.27 ± 4.92% and 90.48 ± 5.95% microbial recovery for human and swine models, respectively, with stability maintained for over 180 days. X-Gutor provides a scalable, modular, and translationally relevant platform for the high-fidelity engineering of gut microbial ecosystems.

**Figure 1.**
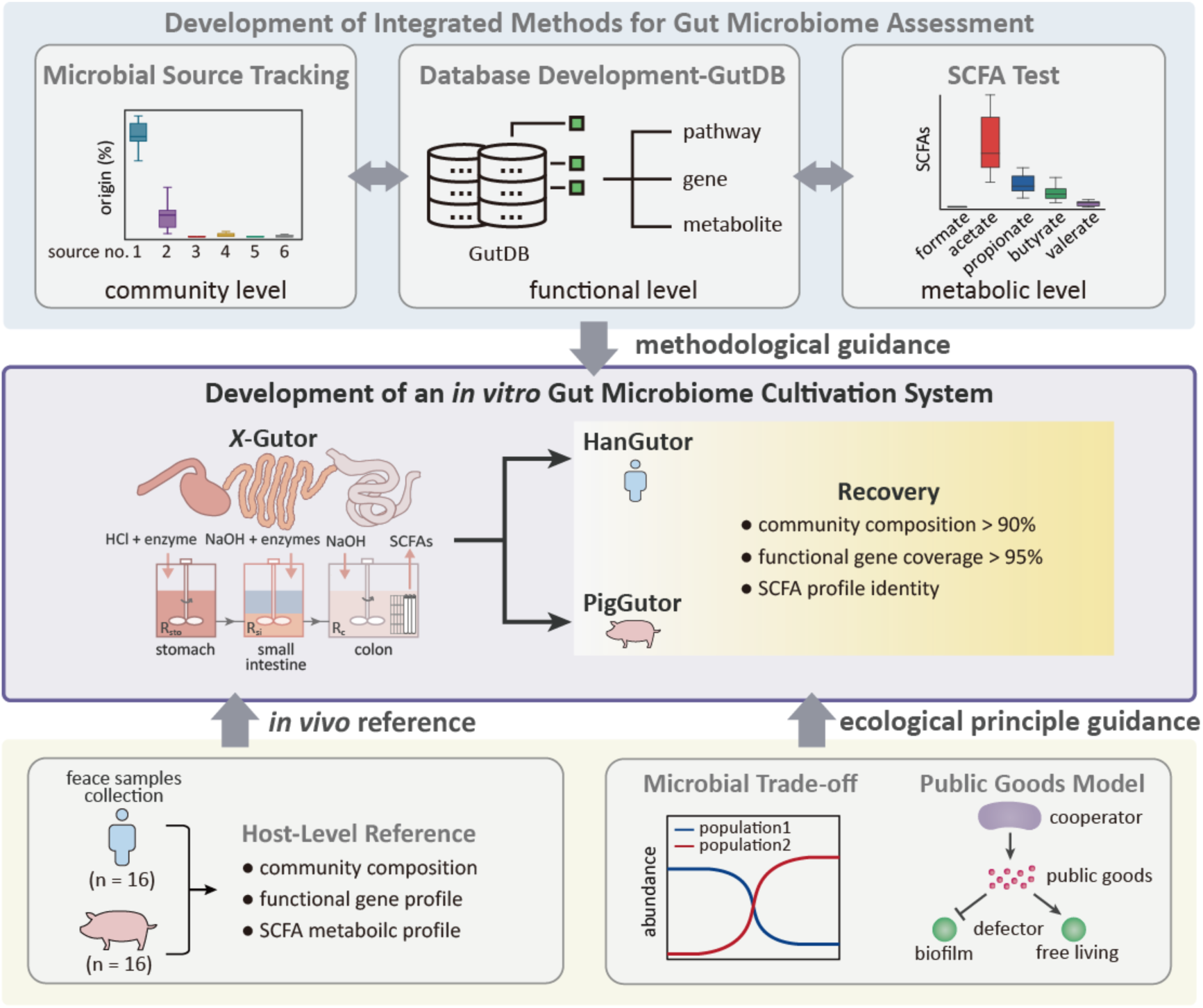
Development of *X*-Gutor platform for mimicking the monogastric digestive system. The system integrates three independent gut-microbiome-profiling methods and referred *in vivo* data from swine and human fecal samples to guide the recreation of *in vitro* gut microbiome in *X*-Gutor.

## Results

### A multi-layer assessment framework to guide the gut microbiota recovery

To guide the recreation of human gut microbiota in the *in vitro* culturing system, we developed an integrated strategy to assess gut microbiota recovery at the taxonomic, functional and metabolic levels. At the taxonomic level, we applied microbial source tracking using the FEAST algorithm, a well-established approach for quantifying the proportional contributions of different source communities to a target community, to resolve the contributions of human and animal gut microbiota to communities established in the *in vitro* culturing system. To encompass the range of physiological and ecological parameters relevant to long-term operation of biosystems that shape the microbial community structure and composition, a total of 1,169 gut microbiota datasets were selected and retrieved from public database (NCBI) as representative source communities. This database comprises poultry (chicken, n=133), monogastric animals (human, n=524; swine, n=270; mouse, n=149), and multigastric animals (cattle, n=93) (Table S1). Together, these hosts represent major axes of gut structure and function, including compartmentalization, hydraulic retention time, dietary substrate complexity and luminal pH, which directly inform biosystem architecture and operating parameters. Principal coordinates analysis (PCoA) of Bray-Curtis dissimilarities revealed strong host-specific clustering of gut microbiota (ANOSIM, R² = 0.962, *P* < 0.001) (Fig. 2A). Samples from each host clustered tightly along the first two principle coordinates, which together captured 41.6% of the total variance, with minimal overlap between different source communities. Consistently, PERMANOVA analysis indicated that host identity explained a substantial fraction of microbiota variation (R² = 0.539, *P* < 0.001), highlighting the driving effects of host-specific factors on shaping gut community structure. To benchmark FEAST’s capacity to resolve host-associated contributions, 50 human gut subsamples were used as a sink for human gut sources, while 50 anaerobic sludge subsamples were included to represent non-gut sources. Source tracking analysis using FEAST revealed that 95.34% (95% CI, 93.56%-96.29%) of the human gut microbial composition was successfully attributed to the intended human sources, while the proportion of unknown sources remained consistently negligible (below 0.12%) (Fig. 2B). In contrast, anaerobic sludge samples were predominantly attributed to unknown sources (87.18%; 95% CI, 84.99%-89.38%), with negligible contribution from human and animal gut microbiota. These results demonstrate the high specificity and accuracy of microbial source tracking using FEAST in quantifying host-specific microbial contributions and provide a rigorous benchmark for evaluating the recovery of gut microbiota reconstruction at the taxonomic level.

**Figure 2.**
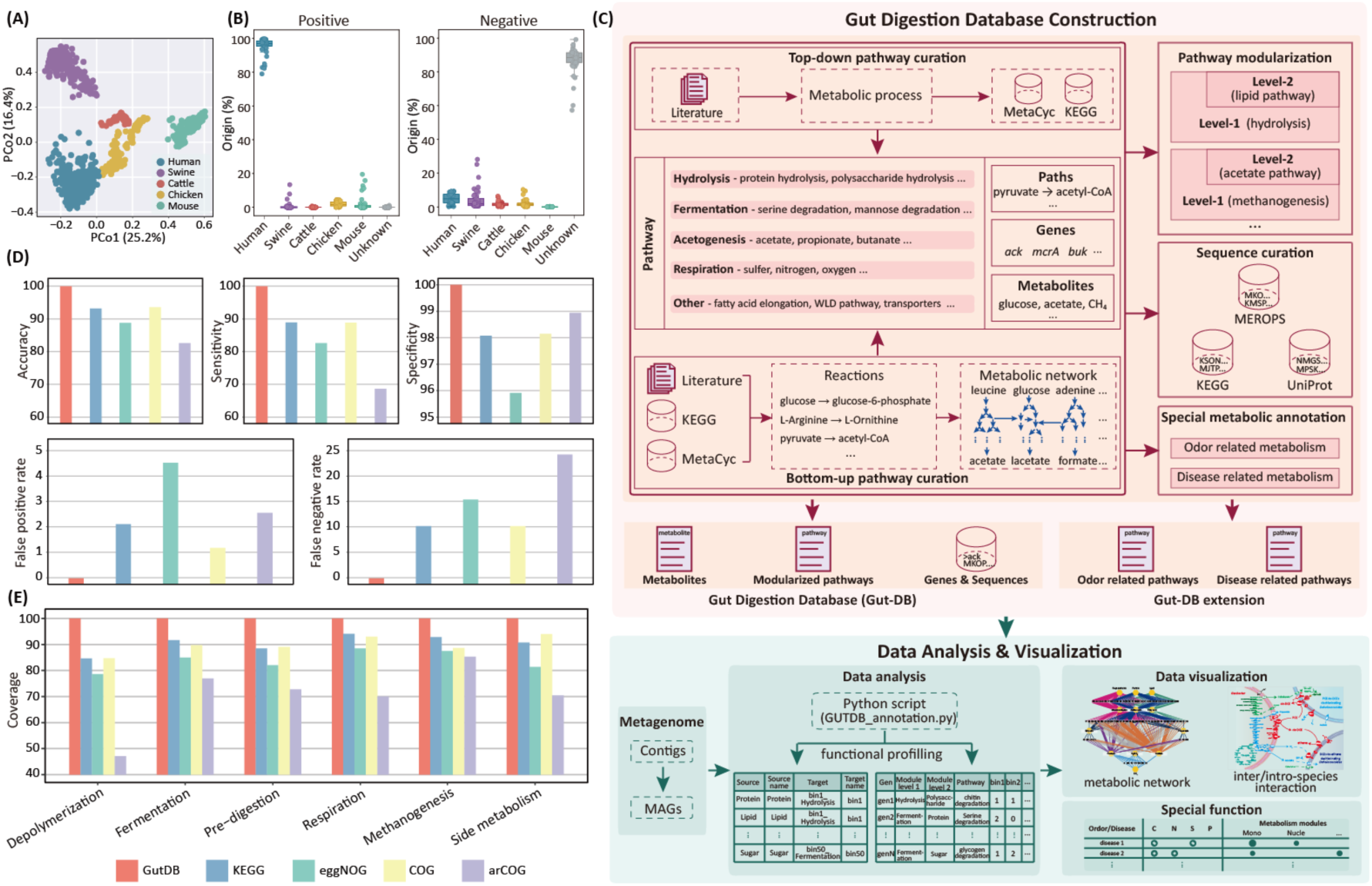
An integrated method for multiple-level assessment of gut microbiota recovery and fiedility. **a**. PCoA of gut communities (n=1,169) from different types of hosts for construction of community sink in microbial source tracking analysis. **b**. Feasibility evaluation of microbial source tracking using FEAST. Positive and negative tests were performed on randomly selected subsets (n = 50) of human gut and anaerobic digestion sludge samples, respectively, to evaluate the specificity and accuracy of the source tracking model. **c.** Process flowchart for construction of the GutDB. Both top-down and bottom-up pathway curation methods were employed to guarantee database accuracy. The curated metabolic pathways, genes, gene-encoded enzymes and metabolites were grouped into metabolism-specific modules, of which sequence data were mainly retrieved from KEGG, UniPort and EMROPS databases. **d** & **e.** The accuracy assessment (including accuracy, sensitivity, specificity, false-negative rate, false-positive rate, and coverage) for KEGG, eggNOG, COG, arCOG and GutDB databases.

Central to the gut microbiota assessment strategy is the curated Gut Metabolism Database (GutDB), which enables high-resolution functional reconstruction and systematic interrogation of gut microbial interactions using meta-omics analyses (Figure 1). The GutDB comprises 7,916 genes and their encoded enzymes, 1,529 metabolites, and 1,890 reactions. These genes, metabolites and reactions are organized into five primary functional modules (level 1) and 16 sub-modules (level 2) that capture key metabolic processes of gut microbiota, including hydrolysis, fermentation, acetogenesis, respiration and other gut-specific functions (Fig. S2). The database was constructed through an integrated top-down and bottom-up strategy (Fig. 2C). In the top-down curation, core gut metabolic pathways were identified from literatures and established resources (e.g., KEGG and MetaCyc), and associated genes, enzymes and metabolites were aggregated into coherent functional modules. Complementing this, the bottom-up approach traced target metabolites to their corresponding genes, enzymes and pathways, enabling the assembly of direct metabolic networks to link substrates with final products. The associated gene and protein sequences were systematically retrieved from KEGG, UniProt, and MEROPS databases to support precise functional annotation. Three specialized modules (Transporter Module, Disease-Related Module, and Odor-Related Module) were incorporated to facilitate studies on gut-associated metabolism and odor production (Fig. 2C).

To evaluate the performance of curated GutDB in metabolic profiling of gut microbiota, an artificial dataset including 10,000 gene sequences from GutDB and 10,000 non-GutDB gene sequences were constructed to compare annotation accuracy, sensitivity, and specificity with KEGG, eggNOG, COG, and arCOG databases. Results showed that GutDB markedly enhanced metabolic annotation accuracy and sensitivity, e.g., 99.98% accuracy and 99.95% sensitivity in metabolic profiling of gut microbita (Fig. 2D). Relative to the widely used KEGG, eggNOG, COG and arCOG databases, GutDB increased annotation accuracy by 6.10-16.56% and sensitivity by 11.02-31.31%, respectively. Notably, arCOG showed substantially lower performance, with an accuracy of 83.42% and a sensitivity of 68.64%, highlighting the limitations of general orthology databases for gut-specific metabolic annotation (Fig. 2D). Module-resolved analysis further revealed that GutDB provided broad functional gains, particularly for depolymerization processes, where annotation accuracy increased by up to 52.94% relative to the arCOG (Fig. 2E).

### Engineering monogastric gut environment in X-Gutor

To establish baseline metabolic and microbial profiles for *in vitro* culturing model development, fecal samples from human (n=16) and swine (n=16) were collected and analyzed for SCFA production (a key metabolic readout of gut microbial activity) and microbial community profiles (Fig. S3; Table S2; Table S3), which offered complementary metabolic and microbial metrics to evaluate how faithfully the *in vitro* system reconstructed native gut communities. SCFA compositions and concentrations were broadly comparable between the two hosts, whereas gut microbial community structures differed markedly between humans and swine (PERMANOVA, R² = 0.492, P < 0.001; Fig. 3A and B). For the swine gut microbiota, microbial profiles were spatially consistent across the three colon regions, i.e. proximal, transverse and distal colon (PERMANOVA, R² = 0.165, P = 0.264; Fig. S4A and B), supporting the employment of a single reactor to mimic the colon. Based on these observations, *X*-Gutor as the *in vitro* gut microbiota culturing system was established to mimic the monogastric digestive system (Fig. 3C), which comprised three sequential reactors representing the stomach (R_sto_), small intestine (R_si_) and colon (R_c_), and were operated under conditions of 37°C with pH control (Fig. 3C).

**Figure 3.**
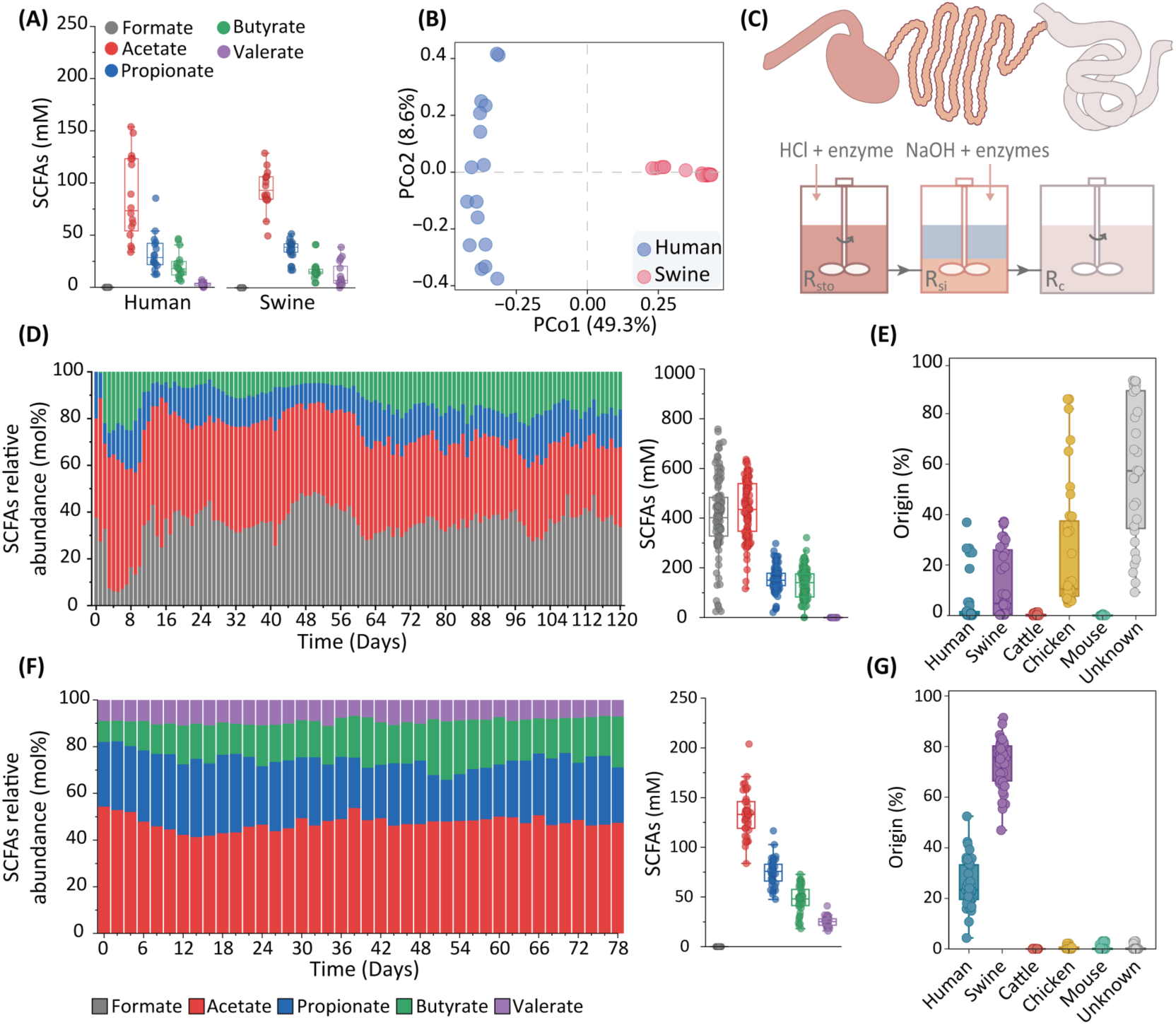
Establishment of *X*-Gutor as an *in vitro* gut microbiome culturing model to mimic human and swine monogastric digestive systems. a. SCFAs composition and concentration in human (n = 16) and swine feces (n = 16). b. Principal coordinate analysis (PCoA) of human and swine feces communities. c. Schematic representation of the *in vitro X*-Gutor model designed to mimic monogastric digestive systems. R_sto_, R_si_ and R_c_ were set to simulate stomach, small intestine and colon, respectively. The intestinal wall simulator was added in the 2^nd^ generation swine colon simulator. d. Concentrations and compositions of SCFAs in the 1^st^ generation swine colon simulator (R_c-P1_). e. Origin estimation of recreated microbiota in R_c-P1_. f. Concentrations and compositions of SCFAs in the 2^nd^ generation swine colon simulator (R_c-P2_). g. Origin estimation of recreated microbiota in R_c-P2_. The origin estimation was based on microbial source tracking analysis.

The *X*-Gutor platform was first configured to mimic the digestive system of swine (designed PigGutor). The hydraulic retention time was set to 1.25 days by regulating influent flow into R_sto_ (stomach simulator) and effluent from R_C_ (colon simulator), while enzymatic digestion in R_sto_ and R_si_ (small intestine simulator) was optimized by adjusting salinity and digestion duration (Fig. S4C; Table S4). In the 1^st^ generation R_c_ (R_c-P1_) of the PigGutor, the metabolic profiles differed significantly from native swine gut samples, e.g., the total SCFA concentration substantially elevated (1133.79 ± 245.14 mM *in vitro vs*. 160.03 ± 37.52 mM *in vivo*; ANOVA, *F* = 320.5, *P* < 0.001), and the SCFA composition dominated by formate (407.93 ± 148.87 mM) that was undetectable *in vivo* (Fig. 3D). The excessive formate production in R_c-P1_ was likely attributed to the use of high concentration of NaHCO_3_ (120 mM) as a pH-buffer, potentially promoting intracellular formate formation via NADPH-dependent formate dehydrogenase activity (Fig. S5A)^26,27^. Importantly, the elevated total SCFAs were not solely attributable to the formate generation since the concentration increase was observed across all major SCFAs. Consistently, microbial source tracking analysis revealed that R_c-P1_ communities were primarily attributed to unknown sources (57.97 ± 28.30%), with a maximum attribution of 38.25% with swine gut microbiota (Fig. 3E; Fig. S5B). Together, these results indicated that R_c-P1_ failed to recreate the swine gut microbiota at both the metabolic and taxonomic levels.

To achieve the high recovery of swine gut microbiota, a 2^nd^ generation swine colon simulator (R_C-P2_) was developed through two major modifications (Fig. S5C; Table S4): (i) replacement of NaHCO_3_ with NaOH for pH control, and (ii) incorporation of a mucin-coated intestinal wall simulator to enable SCFA absorption and biofilm formation. The latter represents a defining feature of *X*-Gutor and consists of a customized 3D-printed polypropylene simulator that replicates the specific structural architecture of the swine or human intestinal wall (Fig. S5C). This structure comprises a rigid frame serving as a biofilm-supporting matrix and hollow fiber membrane to facilitate adsorption of SCFAs and other fermentation products. The frame was coated with mucin type II-agar to promote the establishment of mucosa-associated microbiota, while the hollow fiber membrane was wrapped around the frame to simulate nutrient absorption and biofilm development in the colon (Fig. S5C). These optimizations effectively eliminated formate accumulation and restored SCFA profiles that closely resembled those of native swine gut samples, both in the total concentration (281.10 ± 43.38 mM) and the composition (Fig. 3F). Microbial source tracking analysis demonstrated a marked increase in community similarity between R_c-P2_ and native swine gut microbiota (73.50 ± 9.56%; Fig. 3G; Fig. S5D), indicating substantial improvement in the ecological and functional reconstruction of the swine gut environment.

### Ecological fine-tuning via microbial interaction control

To further improve microbial recovery of swine gut microbiota in the PigGutor, major microbial populations and their correlation with the microbiota recovery ratio in R_c-P2_ were investigated (Fig. 4A and B). While *Succinivibrio* was a major population in R_c-P2_ (Fig. 4A), it represented only a minor fraction of the native swine gut microbiota. Conversely, dominant populations in R_c-P2_, such as *Prevotella*, *Roseburia* and *Megasphaera*, were also prevalent in the native swine gut microbiota (Fig. 4A). Quantitative microbial source tracking (via FEAST) revealed a strong negative correlation between the relative abundance of *Succinivibrio* and the recovery ratio of swine gut microbiota (R^2^ = 0.87, *P* < 0.001; Fig. 4B), as well as a negative correlation between *Succinivibrio* and *Prevotella* (R^2^ = 0.41, *P* < 0.001; Fig. S5D). No significant correlations were observed between the microbiota recovery ratio (or *Prevotella*) and other major populations (Fig. S6). These findings suggested that the *Succinivibrio* may act as a niche competitor to the *Prevotella*, thereby compromising microbial recovery in R_c-P2_. Interestingly, *Succinivibrio* dominated the free-living digestate fraction (24.66 ± 9.37%), yet nearly absent in biofilm communities on the intestinal wall simulator (biofilm matrix) of the same reactor (0.84 ± 0.16%) (Fig. 4A).

**Figure 4.**
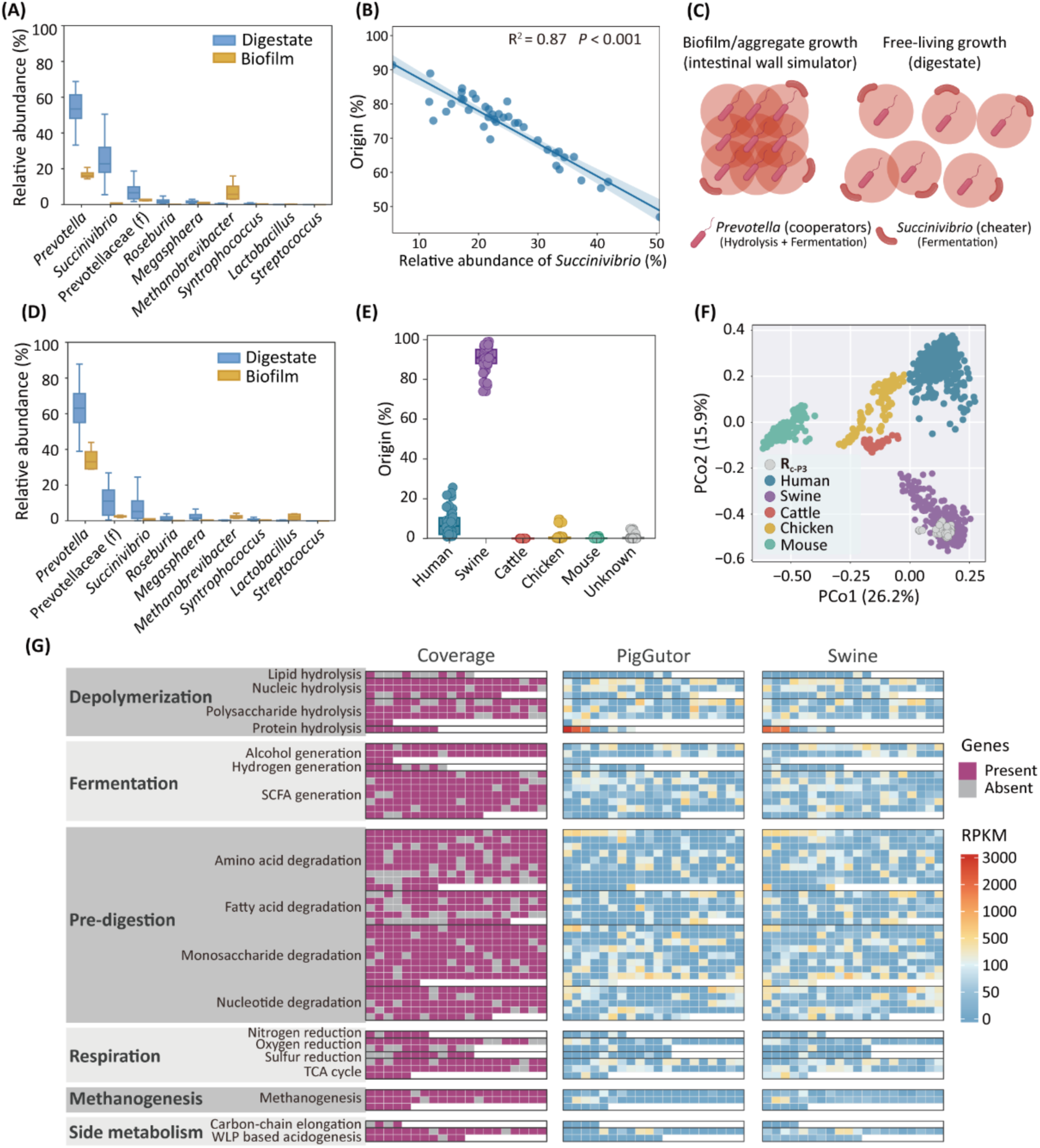
Recreation of swine gut microbiota in PigGutor. **a.** Relative abundance of major populations at the genus level in digestate and biofilm samples of R_c-P2_. **b.** Correlation between relative abundance of *Succinivibrio* and microbial source tracking based recovery ratio of swine gut microbiota in R_c-P2._ *R^2^*, coefficient of determination (n = 40). **c.** Schematic diagram showing the relationship between *Prevotella* (cooperator) and *Succinivibrio* (cheater) in the biofilm and digestate samples based on a spatial ecological public goods model. **d.** Relative abundance of major populations at the genus level in digestate and biofilm samples of R_c-P3_. **e.** Origin estimation of recreated microbiota in R_c-P3_. **f.** PCoA of *in vitro* swine gut communities in R_c-P3_ together with potential source communities. **g.** Coverage and functional profile comparison of the recreated microbiota in R_c-P3_ and *in vivo* swine gut microbiota based on GutDB-supported metagenomic analysis.

The distinctive distribution of *Succinivibrio* in the digestate (fast diffusion niche) and the biofilm (slow diffusion niche) is consistent with a spatial ecological public goods model^28,29^ (Fig. 4C). In this framework, *Succinivibrio,* acting as a “defector” lacking hydrolytic capacity, relies on hydrolytic products (public goods or common resources) released by “cooperators” such as hydrolytic and fermenting microorganisms, including *Prevotella*. In R_c-P2_, the digestate (feces simulator) and biofilm (intestinal wall simulator) represent distinct spatial niches where *Succinivibrio* and *Prevotella* function as a defector (to exploit common resources without contributing) and cooperator (to sustain common resources at some cost), respectively. Defectors exploit common resources without contributing, whereas cooperators sustain these resources at a metabolic cost. Slow diffusion in biofilm enabled cooperators (*Prevotella*) to readily locate and colonize the niche, whereas fast diffusion in digestate promoted coexistence of cooperators and defectors (*Succinivibrio*) as observed in the R_c-P2_ (Fig. 4C).

Based on this ecological insight, the 3^rd^ generation, R_c-P3,_ was designed to suppress the cell growth of *Succinivibrio* by increasing the proportion of biofilm-growth-supporting matrix from 5% (v/v) in R_c-P2_ to 20% (v/v) (Table S4). This adjustment significantly reduced relative abundance of *Succinivibrio* from 24.66 ± 9.37% to 7.06 ± 6.76% (ANOVA, *P* < 0.001; Fig. 4D), while increasing the prevalence of cooperators (e.g., *Prevotella*, uncharacterized Prevotellaceae and *Lactobacillus*) for 78 days operation (Fig. 4D). Consequently, the recovery ratio of swine gut microbiota in the R_c-P3_ increased to 90.48 ± 5.95% (Fig. 4E and F) with an improved SCFAs profile (Fig. S5F). The dominance of *Prevotella* in R_c-P3_ mirrored its natural enrichment in swine gut, likely reflecting the carbohydrates- and fiber-rich diet of swine, consistent with the trends observed in human gut microbiota consuming carbohydrates- and fiber-rich diets^30^. Functional recovery was further assessed with the GutDB-based metagenomic analyses (Fig. 2C; Fig. S2). Results showed that R_c-P3_ captured 95.89% of the major functional traits (with gene abundance ≥5 RPKM) observed in the native swine gut microbiota, with the remaining uncovered genes mainly corresponding to the low-abundant (< 5 RPKM) genes in native gut microbiota (Fig. 4G). In addition to the high microbiota coverage, metabolic profiles of *in vitro* (R_c-P3_) and *in vivo* swine gut microbiota were highly similar (Fig. 4G; Fig. S7A; Table S5), with a strong positive correlation (R² = 0.83, *P* < 0.001) in abundance of functional metabolism modules (Fig. S7B). These results, together with source tracking analyses, demonstrate that R_c-P3_ achieved high levels of taxonomic, functional and metabolic recovery, establishing PigGutor as an effective *in vitro* model for recreating the swine gut microbiome.

### Generalizability to human gut microbiome

To test the capability of X-Gutor in recreating human gut microbiota, an *in vitro* <u>h</u>um<u>an</u> intestine model (HanGutor; Table S4) was developed as a twin gut of feces donor #1 (the person to donate feces for inoculating the human colon simulator R_c-H_), with whom identical food feed was provided (Fig. S8). To better mimic human gut conditions in the HanGutor, digestate retention time (defecation interval) as a key parameter to retain microorganisms with specific cell division and generation time was optimized in batch experiments based on their SCFAs profiles (Fig. S9A; Table S4). In the 1^st^ generation of HanGutor with optimized digestate retention time (2 days), comparable total concentration and composition of SCFAs were generated in R_c-H1_, relative to *in vivo* human gut microbiota (Fig. 5A; Fig. S9B). Nonetheless, only 84.02 ± 9.91% human gut microbiota was recovered in the *in vitro* intestine model (Fig. 5B; Fig. S9C, D). *Bacteroides* as a dominant population in human gut microbiota was shown to positively correlate with microbial recovery ratio of R_c-H1_ (R^2^ = 0.33, P < 0.05; Fig. S9E), in contrast to the predominance of *Prevotella* in PigGutor. In addition, a negative correlation between *Prevotella* and *Bacteroides* was present in the R_c-H1_-recovered human gut microbiota (R^2^ = 0.48, P < 0.01; Fig. S9F). Similar abundance tradeoff between *Prevotella* and *Bacteroides* was observed in human gut microbiota with different diets^31,32^. A closer examination of the major differences between HanGutor and PigGutor suggested that feed composition (diet) and anaerobic condition (redox potential) could be potential factors triggering the *Prevotella*-*Bacteroides* tradeoff. MAGs of *Prevotella* and *Bacteroides* retrieved from recreated microbiota in R_c-H1_ suggested that the *Prevotella* had comparatively less metabolism substrate diversity but higher tolerance to oxidative species (e.g., nitrate reductase and superoxide reductase), relative to the *Bacteroides* (Fig. 5C). Consistently, the 16S rRNA gene-based prediction showed that the high O_2_ tolerance scores were positively correlated with *Prevotella/Bacteroide* ratio in human gut microbiota (R^2^ = 0.55; P < 0.001; Fig. 5D), corroborating the hypothesis of redox potential as a potential *Prevotella*-*Bacteroides*-tradeoff-triggering factor.

**Figure 5.**
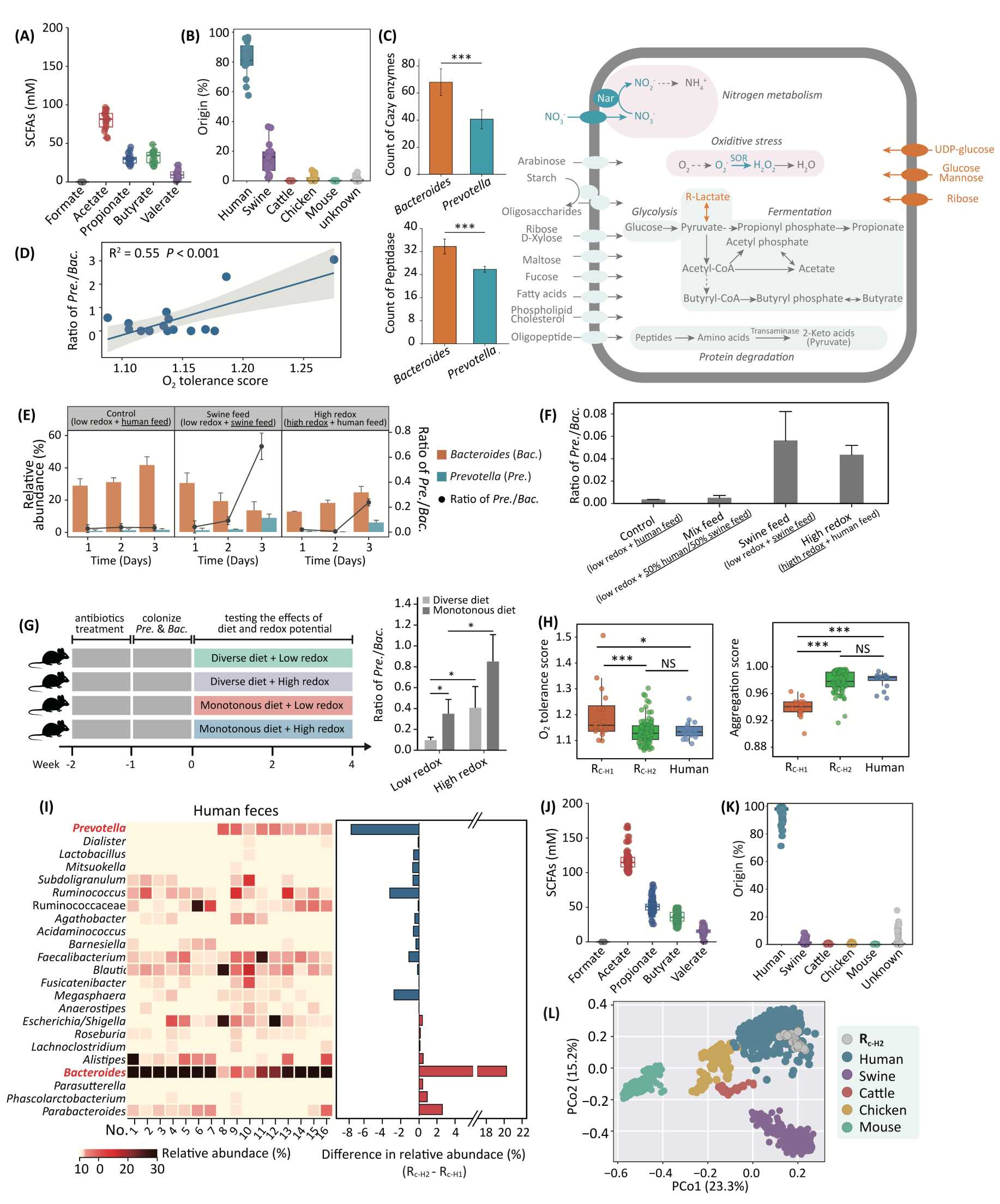
Recreation of human gut microbiota in HanGutor. a. **b.** Origin estimation of SCFAs profile and recreated microbiota in R_c-H1_. **c.** MAGs-based prediction of metabolic potential of *Prevotella* and *Bacteroides in* R_c-H1_. Pathways shared by *Prevotella* and *Bacteroides* were marked in grey arrows; pathways present exclusively in the *Prevotella* and *Bacteroides* were marked in blue and orange arrows, respectively.

To test the effects of diet and anaerobic condition (redox potential) on the human gut microbiota recovery, *in vitro* batch cultivation experiments were setup with a *Bacteroides*-rich fecal sample (Fig. 5E) and *Prevotella/Bacteroides* pure cultures (Fig. 5F). Results showed that increasing redox potential and decreasing diet composition diversity (from human feed to swine feed) increased the relative abundance of *Prevotella* and triggered the *Prevotella*-*Bacteroides* tradeoff (Fig. 5E and F), suggesting their critical roles in selection of *Bacteroides*. The selection of *Bacteroides* over *Prevotella* by low redox potential and diverse diet composition was further confirmed in the *in vivo* experiments using a mice model (Fig. 5G), where a consistent increase of *Prevotella*/*Bacteroides* ratio was observed. Therefore, in the 2^nd^ generation of HanGutor (R_c-H2_), a lower redox potential was implemented as a key control parameter to further improve the gut microbiota recovery (Table S4). In consistence, the O_2_ tolerance score predicted for the microbial communities from R_c-H2_ demonstrated a compatible adaption to lower redox potential to R_c-H2_ but aligning with levels observed in human feces (Fig. 5H). This adaptation is further supported by the aggregation score prediction, which indicated an improved microbial aggregation potential in the R_c-H2_ (Fig. 5H), also consistent with the higher ratio of biofilm supporting matrix in R_c-H2_ (20% in R_c-H1_ *vs.* 30% in R_c-H2_) (Table S3). With the modification implemented in R_c-H2_, 16S rRNA gene-based microbial community profiles showed a relative abundance decrease in *Prevotella* and an increase in *Bacteroides* in R_c-H2_, relative to the R_c-H1_ (Fig. 5I, Fig. S9G). Importantly, these modifications preserved SCFAs profiles comparable to those of native human fecal samples (Fig. 5J; Fig. S9H). Consequently, the recreated microbiota in the 2^nd^ generation *in vitro* intestine model shared a remarkable 96.27 ± 4.92% similarity with *in vivo* human gut microbiota (Fig. 5K and L), corroborating the importance of redox potential in mediation of *Prevotella*-*Bacteroides* tradeoff in human gut microbiota.

Importantly, the R_c-H2_ model demonstrated not only high microbiota recovery in both the microbial community compositions and SCFAs profiles, but also long-term stability (180 days’ stable operation) in the HanGutor (Fig. 5J; Fig. S9H). Metagenomic analyses showed 97.83% coverage of major human gut metabolism traits (with gene abundance ≥5 RPKM) in the R_c-H2_ (Fig. S10A). The overall metabolism module profiles in microbiota of HanGutor (R_c-H2_) and human gut were highly similar (R² = 0.66, *P* < 0.001; Fig. S10B and C; Table S5). Together, these findings demonstrate that the HanGutor (R_c-H2_) can accurately and stably reproduce the structure and function of human gut microbiota. Its tunable operation parameters and scalability provide an ethical and robust platform for investigating gut microbiota responses to diet, intestinal physiological or pathological conditions, and probiotic interventions.

Data on the number of Cazy enzymes and peptidases were presented as means ± s.e.m. Nar, nitrate reductase; SOR, superoxide reductase. **d.** Correlation between 16S rRNA gene-based prediction of O_2_ tolerance and *Prevotella/Bacteroides* ratio in native human gut microbiota. The O_2_ tolerance score ranged from 0 to 5, representing obligate anaerobic to obligate aerobic conditions. Change in relative abundance and ratio of *Prevotella* (*Pre.*) and *Bacteroides* (*Bac*.) in batch experiments setup with fecal samples **(e)** or Prevotella-and-Bacteroides pure cultures **(f)** and with low/high (obligate-anaerobic/microaerobic) redox potential and human/swine feeding substrates. **g.** Change in relative abundance ratio of *Prevotella* (*Pre*.) and *Bacteroides* (*Bac*.) in the 4-week *in vivo* experiments with mice fed with diverse/monotonous diet under low and high redox potentials. **h.** 16S rRNA gene-based prediction of O_2_ tolerance and aggregation potential of R_c-H1_, R_c-H2_, and human gut microbiota. The O_2_ tolerance score ranged from 0 to 5, representing obligate anaerobic to obligate aerobic conditions, while aggregation score ranged from 0 to 1, stranding for no aggregation to observed aggregation. **i.** Major microbial populations at the genus level in human feces and their difference in relative abundance between R_c-H1_ and R_c-H2_. **j. k.** Origin estimation of SCFAs profile and recreated microbiota in R_c-H2_. **l.** PCoA of *in vitro* human gut communities in R_c-H2_ together with potential source communities. Statistical significance was calculated based on ANOVA, and denoted with asterisks (*), \**P* < 0.05, \*\*\**P* < 0.001.

## Discussion

The *in vitro* gut microbiome culturing model offers not only an economic-effective and ethically favorable alternative to animal models but also a customizable and host-independent platform for interrogating gut microbiota ecology, host-microbe interactions, and intestinal health^7,9^. Nevertheless, their biological relevance of existing systems has been fundamentally limited by the challenge of achieving both high microbiota recovery and long-term ecological stability, two benchmarks for faithful reconstruction of native gut system^7,9,17^. In this study, we establish X-Gutor as an *in vitro* gut microbiome culturing model that overcomes these limitations by sustaining microbial community composition and gene-encoded functional properties highly comparable to native host gut microbiota over extended periods without repeated microbial supplementation, representing a substantial advance in the faithful reconstruction of complex gut ecosystems. Beyond its core culturing capability, the *X*-Gutor exhibits a versatile and translationally relevant experimental platform: (1) as a customizable and scalable platform for probiotics and drug development, as well as for systemic evaluation of chemical exposures and their potential risks for gut microbiota and the host health; (2) as a biosystem for expanding stool-bank-derived healthy gut microbiota for fecal microbiota transplantation, alleviating donor-source limitations and enabling cultivation of previously uncharacterized gut “microbial dark matter”; (3) as a framework to guide gut-microbiome-oriented personalized nutrition and other microbiota-targeted interventions; and (4) as a modular system that can be integrated with host cell cultures to facilitate mechanistic elucidation of host-microbiome interactions under controlled conditions.

To ensure sustained microbial community reproducibility and biological relevance, we implemented a data-driven, multi-layered evaluation framework that integrates microbial source tracking, GutDB-enabled meta-omics, and short-chain fatty acid profiling as complementary and interdependent assessment layers. Together, these metrics capture taxonomic composition, gene-encoded functional potential, and metabolic output, enabling quantitative benchmarking of *X*-Gutor cultures against native host gut ecosystems and supporting iterative, evidence-based optimization of system parameters. In particular, the curated metabolic architecture of GutDB permits high-resolution and computationally efficient reconstruction of gut-specific metabolic functions, thereby addressing a persistent disconnect between community structure and ecological function in *in vitro* models. By anchoring functional inference in experimentally validated gut metabolic pathways, this framework strengthens the causal interpretation of reconstructed microbial functions. Importantly, the modular nature of the *X*-Gutor system and its evaluation pipeline confers adaptability beyond monogastric systems, providing a generalizable framework for the rational design, validation, and cross-host comparison of customized *in vitro* microbiome models. As such, this integrative assessment strategy establishes a scalable paradigm for advancing *in vitro* gut ecosystem modeling from descriptive community maintenance toward mechanistically informed and translationally robust platforms.

*X*-Gutor directly addresses the limitations of murine models in mechanistic gut microbiome research, which is often constrained with limited microbial community overlap with humans and poor translatability to human physiology and disease ^7,33^. By implementing a systemic and ecology-informed optimization strategy, *X*-Gutor enables high-reproducibility reconstruction of host gut microbiota. The process begins with “coarse tuning,” achieved exclusively through the manipulation of physicochemical and operational reactor conditions, resulting in a microbiota recovery rate of 73.50 ± 9.56%, thereby demonstrating that substantial community reproducibility can be attained without exogenous microbial supplementation. These design parameters, including influent composition, inoculum characteristics, biomass retention time, and a specialized 3D-printed hollow fiber architecture that mimics the structure of the intestinal wall, as well as niche-defining factors such as redox potential, temperature, and pH, closely parallel the primary determinants of gut microbiota assembly *in vivo*, establishing a mechanistic link between engineered reactors and host-associated ecosystems. Following this, “fine tuning,” involved targeted analysis of dominant species interactions, specifically *Prevotella* and *Succinivibrio* in PigGutor, and *Prevotella* and *Bacteroides* in HanGutor, further enhancing microbiota recovery to over 96% and restoring short-chain fatty acid profiles to align with those of native host microbiota, confirming both compositional and functional congruence. This comprehensive approach positions *X*-Gutor as a versatile framework for reconstructing diverse gut ecosystems with high structural and metabolic fidelity, while also providing a generalizable methodology that can be applied to various *in vitro* gut models, transcending host and species limitations.

In contrast to a relatively complete atlas of gut microbiota taxa, limited progress has been made in elucidating the complicated web of interactions of the gut microorganisms^1,34^. *Prevotella* as a major population present in both human and swine gut microbiota plays a key role in gut homeostasis through forming complicated interactions with other microorganisms, including our observations of: (i) a diet and redox-potential regulated abundance tradeoff between *Prevotella* and *Bacteroides*, and (ii) cooperator-defector-like competition for public goods between *Prevotella* and *Succinivibrio*. More importantly, key factors regulating the interactions of *Prevotella* with *Bacteroides* and *Succinivibrio* are diet/redox-potential and biofilm-supporting-matrix, respectively. Insight into these key interaction-regulating factors enable development of new strategies to remediate gut microbiota dysbiosis. For the tradeoff between *Prevotella* and *Bacteroides*, diet has long been believed as the key driving factor based on large-scale survey metadata^31,32,35^, but studies in humans on short-term dietary change showed the triggering effects of yet-to-be-characterized factors^36^. In our study, both HanGutor and batch experiments showed enhanced growth of *Bacteroides* and its tradeoff with *Prevotella* upon decreasing oxygen exposure and consequently lowering redox potential. The *Bacteroides* generally occurred in obligate anaerobic environments with extremely low redox potential including methanogenic bioreactors and microbial reductive dechlorination consortium^37,38^, whereas *Prevotella* could coexist with various facultative or oxygen-tolerant anaerobes and predominate anoxia niches, e.g. human skin, mouth and vagina^30,39^. Numerous previous studies have shown that the increased relative abundance of *Prevotella* could be associated with varied intestinal dysbiosis and consequent mucosal inflammation ^40,41^, which was generally accompanied with available oxygen and nitrate in the colon due to impairment of host control of their availability^35^. In contrast, conditions with high availability of host-derived oxygen and nitrate (high redox potential) in colonic lumen cause dysbiosis by fueling growth of facultatively anaerobic bacteria in colonic microbiota^35,42–44^. Therefore, redox potential, in addition to diet, could be a key factor in triggering the *Prevotella*-*Bacteroides* tradeoff that might further result in *Prevotella*-associated gut dysbiosis. In addition, the newly developed *X*-Gutor and associated method could provide powerful tools for future in-depth studies on the *Prevotella*-*Bacteroides* tradeoff and other microbial interactions, which will facilitate more comprehensive understanding of mechanisms driving the change of gut microbiota and enable exploring the effects of dietary factors, specific intestinal physiological/ pathological conditions, and probiotic use on gut microbiome.

## Methods

### Collection and characterization of feces samples

Human (n=16) and swine (n=16) fecal samples , as well as digesta from swine proximal, transverse and distal colons (n=5/each section, total n=15), were collected and processed according to standard protocols as described^45^ for physiochemical characterizations, microbial community profiling and inoculation of *in vitro* models. Human feces samples were collected according to experimental procedures approved by the Institutional Feces Collection and Use Committee of the Sixth Affiliated Hospital, Sun Yat-sen University. The swine feces and digestion samples of swine proximal, transverse and distal colons were collected from a swine farm in the Guangdong Academy of Agricultural Sciences (Guangzhou, China). The swine feces and digestion samples were collected according to experimental procedures approved by the Laboratory Animal Ethics Committee of Institute of Animal Science, Guangdong Academy of Agricultural Sciences. Details of these samples were provided in Table S2 and S3. The swine intestines used in this study were obtained as by-products of regular food processing.

### Reactor setup and operation (X-Gutor: PigGutor, HanGutor)

#### X-Gutor platform architecture

The X-Gutor consists of three sequential stirred-tank reactors to simulate niche conditions of the stomach (R_sto_, 800 mL), small intestine (R_si,_ 1000 mL) and colon (R_c_, 800 mL), maintained at 37 °C in a water bath. All reactors were continuously sparged with N_2_ to maintain anaerobiosis. The hydraulic retention time (HRT) was set to 1.25 days for PigGutor and 2 days for HanGutor by adjusting daily influent and effluent flow.

#### Enzymatic digestion in R_sto_ and R_si_

Only cell-free enzymatic processes were stimulated in R_sto_ and R_si_ by adding hydrolytic enzymes (Macklin; Shanghai, China), i.e., 738 U/ml pepsin in the R_sto_ and 221 U/ml amylase, 69 U/ml trypsin, 9 U/ml chymotrypsin and 3 U/ml lipase in the R_si_ (Table S3), based on the facts that amylase is mainly secreted by the salivary glands, while pepsin, trypsin, chymotrypsin and lipase are mainly derived from the pancreas.

#### Colon simulator with intestinal wall mimic

The colon reactor R_c_ contained a custom 3D-printed polypropylene frame and wrapped with hollow fiber membrane to simulate nutrient absorption and biofilm support, designed using SolidWorks 2019 and manufactured with a 3D printer (BlueMaker3d, LOBOTICS; Shanghai, China). The frame was coated with mucin type II-agar to support the growth of mucosa-associated microbiota. The hollow fiber membrane was winded around the frame structure to mimic nutrient absorption and biofilm formation in colon (Fig. S5C). The volume ratio of biofilm matrix to digestate was adjusted from 5% (R_c-P2_) to 20% (R_c-P3_) for PigGutor, and from 20% (R_c-H1_) to 30% (R_c-H2_) for HanGutor.

#### Inoculation and operation

Fresh feces (200 g) from a single human donor (#1) or from a pooled swine farm sample were homogenized in 500 mL deionized water and mixed 1:1 (v/v) with the predigested feed from R_si_. This mixture was used as inoculum for R_c_. Reactors were operated in sequencing batch mode with stirring at 100 rpm. pH was controlled by automated addition of 2 M NaOH (instead of NaHCO₃ to avoid formate accumulation). Identical feeding substrates were provided for the PigGutor with that of the swine farm in the Guangdong Academy of Agricultural Sciences, i.e., 50% corn, 20% soybean, 20% chicken powder, 5% bran and 5% feed premix (w/w). Influent substrates of the *in vitro* human gut microbiome culturing model (HanGutor) were identical with the daily diet of feces donor #1 (feces donor for inoculation of the HanGutor) as shown in Fig. S8, i.e., around 22% meat, 38% vegetables, 25% rice and 15% fruit (w/w). Redox potential was measured online and maintained. Detailed information on the influent composition and reactor operation parameters were summarized in Table S4.

#### Batch experiments

To test redox potential and diet effects on *Prevotella*/*Bacteroides* trade-off, batch cultures were setup in serum bottles with *Prevotella copri* DSM18205 and *Bacteroides uniformis* ATCC 8492 pure cultures (Mingzhou Biotechnology Co., Ltd.; Ningbo, China) or with *Bacteroides*-dominated fecal samples (from feces donor #1). Conditions included redox potential (microaerobic and obligate anaerobic conditions) and feeding substrates (i.e., 100% swine feed, 50% swine feed + 50% human daily diet, 100% human daily diet). All batch experiments were run in triplicate and transferred three times before analysis. The 16S rRNA gene copies of *Prevotella* and *Bacteroides* were quantified using species-specific qPCR primer sets Pre51F/Pre248R (5’-CAAGTCGAGGGGAAACGACA-3’/5 -GCTAATCAGACGCATCCCCA-3’) and Bac620F/Bac838R (5’-CGTAAAATTGCAGTTGAT-3’/5’-ATATCGCAAACAGCGAGTA-3’), respectively.

### Chemical analyses

Concentrations of short chain volatile fatty acids (SCFAs) were measured using the method as previously described^46^. Briefly, samples were thawed, weighed (1 g each), acidified with m-phosphoric acid (6.25%, w/w), sonicated and stored overnight at -20°C in refrigerator. Samples were then thawed and centrifuged for 10 min at 16,500× g, and the supernatant was collected for analysis using an Agilent 7890B gas chromatography (GC) equipped with a flame ionization detector (FID; Agilent, Wilmington, DE, USA) and a DB-FFAP column (30 m × 0.25 mm × 0.25 μm; Agilent). Amplicon sequencing, data processing and microbial source tracking-based origin estimation

### Amplicon sequencing and data processing

Genomic DNA (gDNA) for 16S rRNA gene amplicon sequencing was extracted with TINAamp soil DNA Kit (Tiangen, Beijing, China) following the manufacturer’s instructions. The V4-V5 regions of the 16S rRNA genes were amplified with a U515F/U909R primer set, and the amplicons were purified as described previously^47^. To multiplex the PCR amplicons, a specific 8-mer barcode was added to the forward and reverse primers. The purified PCR products were pooled and sequenced on an Illumina Novaseq platform (PE250; Illumina; San Diego, CA, USA) provided by AZENTA (Suzhou, China). The provided paired-end (2×250 bp) reads were filtered, dereplicated, merged, chimera identified and inferred to amplicon sequence variants (ASVs) by using the DADA2 package (version 1.6)^48^ in R software (version 4.1.2). The taxonomy was assigned by using RDP naïve Bayesian classifier^49^ with the SILVA v132 database^50^. To minimize the bias of sequencing depth, ASVs table was rarefied to 21,900 sequences per sample. The principal coordinates analysis (PCoA) based on Bray-Curtis dissimilarity matrix was performed at the ASV level to reveal the community difference.

### Microbial source tracking (FEAST)

Microbial source tracking analysis based on a FEAST method^51^ was used to estimate the origin of targeting microbiota (sink community) recreated in the *in vitro* models, and a highly efficient expectation-maximization-based method was applied to assess the contribution of source communities to a sink community as described^52^. The microbial source tracking method can be act as a metric of community similarity considering both presence/absence of specific microbial populations and their relative abundance ranking order in the sink community^51^. Briefly, 1,169 sets of gut microbiota data were collected as representatives of birds (chicken), monogastric (human, swine and mouse) and multigastric (cattle) digestive systems (Table S1). These gut microbiota data as source communities were first retrieved from NCBI’s Sequence Read Archive (SRA) database, and divided into groups according to sample sources, followed by processing using DADA2 (v3.16)^48^ to obtain the ASV table. To ensure that the selected gut communities from different sources could reflect their respective microbiota, samples were subjected to principal coordinates analysis (PCoA; Fig. 2A) and permutational multivariate analysis of variance (PERMANOVA) using the vegan R package^53^. To evaluate the feasibility of using FEAST method to quantify the contribution of varied source communities to simulated gut microbiota, 50 subsamples were randomly selected from the collected human gut community data as a positive test set and another 50 subsamples from anaerobic sludge digestion microbiota as a negative test set, of which origin evaluation results showed high resolution of the FEAST method-based source tracking analysis (Fig. 2B). The relationship between source and sink communities was finally illustrated using FEAST package in R^51^.

### Traits prediction of gut microbiota

The potential of microbiota to tolerate O_2_ and to form aggregate were predicted using a protocol as described^54^. In brief, a phylogeny that contained taxa from gut or *X*-Gutor samples and LTP taxa^55^ with formally described type specimens and Latin binomials was constructed. The trait data curated from literature and online repositories were mapped onto the tips of the phylogeny using the Latin binomials. The unknown trait values were inferred by using hidden state prediction. The O_2_ tolerance score ranged from 0 to 5, and increasing score indicates the trend from obligate anaerobic to aerobic conditions. The aggregation score ranged from 0 to 1, meaning never observed aggregation to observed aggregation. Analysis of variance (ANOVA) was used to evaluate the significance of difference between the compared groups.

### Establish of gut metabolism database (GutDB)

Establishment of GutDB was detailed in result section. To evaluate the accuracy of GutDB, an artificial dataset including 10,000 GutDB gene sequences and 10,000 non-GutDB gene sequences was constructed to compare the accuracy, sensitivity, specificity, false-positive rate, false-negative rate, and coverage of GutDB with eggNOG, KEGG, COG, and arCOG. The dataset was searched against GutDB, KEGG, COG, and arCOG using USEARCH (identity >30%), and against the eggNOG database using eggNOG-mapper (e-value of ≤1e-4).

### Mice model and microbial colonization of mice gut microbiome

Healthy C57BL/6J male mice were purchased from the Laboratory Animal Center of Sun Yat-sen University and housed in specific pathogen-free (SPF) animal facility in individually ventilated cages with a maximum of four mice per cage. All of the mice used in this study were maintained with free access to food and water under controlled conditions with 22±2°C, 60-70% humidity, and 12-h-dark/light cycle. To study the impact of diet on mice, the colonized mice were divided into two groups and fed for 4 weeks with a customized diverse diet (major nutrients include 27.3% fat, 20.5% protein and 52.2% carbohydrate, w/w) or a monotonous diet (major nutrients include 0% fat, 0.3% protein and 99.7% carbohydrate, w/w) (Table S6), being amended with the necessary trace elements and supplements to maintain normal physiological function in mice. To test the effects of intestinal redox potential on mice, the colonized mice were anesthetized and the anus was then enlarged with 3 mm diameter hoses for 4 weeks to increase the intestinal redox potential of the mice. 200 mg feces from each mouse were collected for DNA extraction and 16S rRNA gene quantification. All animal experiments were reviewed and approved by the Laboratory Animal Care and Use Committee of Sun Yat-sen University and the Laboratory Animal Ethics Committee of Sun Yat-sen University under registration SYSU-IACUC-2023-001868.

*Prevotella copri* DSM18205 and *Bacteroides uniformis* ATCC 8492 were colonized in mouse intestine with different redox potentials, feeding conditions *in vivo*. Prior to colonization, mice were gavaged three times a week for one week with 200 μL antibiotics cocktail (0.1 g/L Ampicillin, 1 g/L Metronidazole, 1 g/L Neomycin trisulfate, and 0.5 g/L Vancomycin) to deplete the intestinal microbiota. To colonize *Prevotella copri* DSM18205 and *Bacteroides uniformis* ATCC 8492 in mouse intestine, these pure cultures or their co-cultures were centrifuged and resuspended in sterile PBS to prepare a mixed bacterial suspension containing 10^9^ CFU of each species, and then were gavaged into the antibiotic-treated mice three times a week for one week.

### Metagenomic analyses

Feces and digestates samples for metagenomic analysis were collected from human (feces donor #1 in Table S1), swine, PigGutor (R_c-P3_, day 58), and HanGutor (R_c-H2_, day 119; R_c-H2-1_ day 80; R_c-H2-2_ day 80). The quality and quantity of the extracted gDNA were evaluated with gel electrophoresis and a Quantus fluorometer (Promega; Madison, WI, USA). The library preparation and metagenomic sequencing on a HiSeq platform (Illumina) were conducted in BGI (Shenzhen, China). Metagenomic sequencing raw data were filtered to remove low-quality bases/reads using trim_galore (v0.6.10) with default parameters. Each metagenome was assembled using metaSPAdes v 3.10.0 (k-mer: 33, 77, 99)^56^. Genome reconstruction of the gut/*X*-Gutor microbe was performed with the function modules of metaWRAP (v1.2)^57^. In brief, contigs with lengths of >1000 bp were used for genome binning. Metagenome-assembled genomes (MAGs) were constructed with contigs from each sample using three different binning algorithms of ‘--metabat2 –maxbin2 --concoct’ in the metaWRAP software. Refinement of bins was performed using the bin_refinement module of metaWRAP. To obtain optimal genome quality, metagenomic sequencing reads were mapped to each bin, and then reassembled with metaSPAdes via the reassemble_bins module of metaWRAP. The completeness and contamination of each bin were evaluated using CheckM^58^. dRep v3.4.3^59^ was used to dereplicate MAGs at 95% ANI. Protein coding sequences (CDS) of metagenomic contigs were determined using Prodigal (v2.6.3)^60^. Functional annotations were conducted based on GutDB using DIAMOND^61^ with an E value threshold of “1e-5”. Potential metabolic modules in metagenome were classified according to GutDB database. To reconstruct the metabolic pathways, the predicted genes of MAGs were further annotated using KEGG automatic annotation server (KAAS)^62^ and dbCAN3^63^. GTDB-Tk 2.0.0 with GTDB taxonomy release_207 was used to assign the taxonomy of MAGs^64^.

Abundance of genes and MAGs were estimated using CoverM (v 0.6.0, https://github.com/wwood/CoverM/), which was based on genes/MAGs coverage and normalized as reads per kilobase per million (RPKM). The similarity comparisons of gene relative abundance (RPKM) in major functional modules between the swine/human feces samples and PigGutor/HanGutor samples were calculated using the equation:

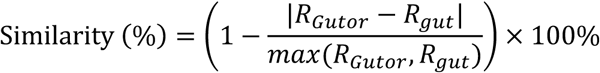

where R_Gutor_ is the total relative abundance (RPKM) of each sub-module in PigGutor/HanGutor samples, and R_gut_ is the total relative abundance (RPKM) of each sub-module in swine/human feces samples.

### Statistical analysis

All batch experiments were carried out at n = 3-5 (see methods and figure captions), and results and error bars indicate mean ± s.d. A one-way ANOVA was applied to test the statistically significant differences between the compared groups using R package stats. *P* values less than 0.05 were considered significant.

## Data availability

The authors declare that the data supporting the findings of this study are available within the article and its supplementary information. Illumina sequencing data for this study have been deposited to the European Nucleotide Archive (ENA) database with an accession number of PRJEB70509. The GutDB can be accessed via the Zenodo repository (https://doi.org/10.5281/zenodo.14837126; Released time was set to 30^th^ December 2026).

## Acknowledgements

This study was financially supported by the Open Funding Project of State Key Laboratory of Microbial Metabolism (Shanghai Jiao Tong University, MMLKF24-06), the Medicine-Engineering Convergence Seed Fund (2025) at Sun Yat-sen University and the National Natural Science Foundation of China (42161160306 and 81600419).

## Author information

These authors contributed equally: Qihao Li, Siyan Zhao.

## Contributions

S. W. designed the study. J. S. and Y. W. carried out *X*-Gutor setup and operation. J. S. and Z. L. established the GutDB database. Q. L. and X. W. conducted mice experiments. Q. L., S. Z. and Z. L. analyzed the data and performed data visualization. J. H. and X. M. helped to collect the human and swine feces samples, respectively. S. Z., S. W. and L. W. drafted the manuscript with the help from X. Z., J. Y., G. S., Z. H. All authors reviewed the results and approved the final version.

## Ethics declarations

The authors declare no competing interests.

## Supplementary information

### Supplementary table legends

**Table S1.** Accession numbers of previously sequencing data for microbial source tracking analysis.

**Table S2.** Basic information of human feces donors.

**Table S3.** Basic information of swine digestate/feces donors.

**Table S4.** Parameters for setup and operation of the *in vitro* gut microbiome culturing models.

**Table S5.** ANOVA test on the functional gene abundances in metabolism modules between PigGutor and swine feces, between HanGutor and human feces.

**Table S6.** Mice diet compositions.

## Supplementary figure legends

**Figure S1. Origin estimation of recreated microbiota in the SHIME system** (**a**) and gut-on-chips (**b**) based on microbial source tracking analysis.

**Figure S2. General and special metabolism modules in the GutDB.** The curated metabolic pathways, genes, gene-encoded enzymes and metabolites were grouped into 5 metabolism-specific modules including organic hydrolysis, fermentation, acetogenesis, respiration, methanogenesis, and gut-microbiota-specific function modules. The modules were numbered with pound sign (#).

**Figure S3. Schematic diagram of the experiment.** The experiments include four parts: (1) community and SCFAs analyses of feces samples collected from swine (n = 16) and human (n = 16) donors, and the data were referred to guide the recreation of *in vitro* gut microbiome; (2) development of three methods (i.e., microbial source tracking, Gut-DB based meta-omics and SCFAs analyses) to support assessment of gut microbiome recovery at the taxonomy, gene-coded function and metabolism levels, respectively; (3) development of the *X*-Gutor to mimic monogastric digestive systems that was further extended to construction of PigGutor and HanGutor; (4) batch experiments based on both pure and mixed cultures were used to test the impact of redox potential and feed on the gut microbiome, particularly the microbial interactions. **Figure S4**. **Microbial and SCFA characterization of swine gut microbiota. a.** SCFAs concentration and profile of swine colon (proximal, transverse and distal colon) samples. **b.** Principal coordinate analysis (PCoA) of the proximal, transverse and distal colon communities. **c**. Impact of high/low salinity and digestion time on digestion efficiency quantified with dissolved organic matter in stomach (R_sto_) and small intestine (R_si_) simulators. Feeding substrates were first digested in the R_sto_ for 10 hours (Stage-I, 0 – 10 hours) and then in the R_si_ during subsequent 38 hours (Stage-II, 10 – 48 hours). In each stage, 3 sets of experiments were setup. In Stage-I: Salt-L-Control, control experimental set without pepsase addition and in low salt condition; Salt-L-Ezn_P_, experimental set with pepsase addition and in low salt condition; Salt-H-Ezn_P_, experimental set with pepsase addition and in high salt condition. In Stage-II: Salt-L-Ezn_P_-Control, control experimental set conducted with the Salt-L-Ezn_P_ digestion product (from Stage-I) without addition of mixed enzymes; Salt-L-Ezn_P_-Ezn_mix_, experimental set conducted with the Salt-L-Ezn_P_ digestion product with mixed enzymes amendment; Salt-H-Ezn_P_-Ezn_mix_, experimental set conducted with the Salt-H-Ezn_P_ digestion product with mixed enzymes addition. See detailed information on salts and digestion enzymes, as well as their concentrations, in **Table S4**.

**Figure S5. Recovery improvement of *in vitro* swine gut microbiota in PigGutor. a**. Formation of formate from CO_2_ and NADPH under mediation of NADP-dependent formate dehydrogenase. PCoA of *in vitro* swine gut communities in R_c-P1_ (**b**), and R_c-P2_ (**d**), together with potential source communities. **c**. Correlation between relative abundance of *Succinivibrio* and *Prevotella* (n = 40). **e**. Optimization of PigGutor by replacing NaHCO_3_ with NaOH as a pH adjustment chemical and by adding mucin-coated intestinal wall simulator. **f**. Temporal change in SCFAs concentration and composition in the 3^rd^ generation swine colon simulator (R_c-P3_).

**Figure S6. Correlations of major populations’ relative abundance with microbial source tracking-based recovery ratio of swine gut microbiota in R_c-P2_ and with the relative abundance of *Prevotella.*** Correlations between the relative abundance of *Roseburia* (**a**), *Megasphaera* (**c**), and *Methanobrevibacter* (**e**) and the recovery ratio of swine gut microbiota in R_c-P2_. R^2^, coefficient of determination (n = 40). Correlations between the relative abundance of *Roseburia* (b), *Megasphaera* (**d**), and *Methanobrevibacter* (**f**) with the relative abundance of *Prevotella* (n = 40).

**Figure S7. Similarity comparison of metabolism modules between *in vitro* and *in vivo* swine gut microbiota based on the GutDB-supporting metagenomic analyses. a**. Similarity of major metabolism modules between the swine gut microbiota and the recreated microbiota in R_c-P3_. **b**. Correlation of functional gene abundance between swine gut microbiota and recreated microbiota in R_c-P3_.

**Figure S8. Foods used as feedstock for the HanGutor.**

**Figure S9. Characterization of recreated human gut microbiota in HanGutor. a**. Impact of digestate retention time on SCFAs generation. **b**. Temporal changes of SCFAs concentration and composition in R_c-H1_. **c**. SCFAs generation in R_c-H1_. **d**. PCoA of *in vitro* human gut communities in R_c-H1_, together with potential source communities. **e**. Correlation between the relative abundance of *Bacteroides* and the microbial-source-tracking-calculated recovery ratio of recreated microbiota in R_c-H1_ (n=16). **f.** Correlation between the relative abundance of *Bacteroides* and *Prevotella* in R_c-H1_ (n=16). **g**. Temporal change in major populations (average relative abundance > 0.1%) at the genus level in R_c-H1_. **h**. Temporal change in major populations (average relative abundance > 0.1%) at the genus level in R_c-H2_. **i**. Temporal changes of SCFAs concentration and composition in R_c-H2_.

**Figure S10. Comparison of *in vitro* and *in vivo* human gut microbiota at the functional level. a**. Coverage and functional profile comparison of the recreated microbiota in R_c-H2_ and *in vivo* human gut microbiota based on GutDB-supported metagenomic analysis. **b**. Similarity of major metabolism modules between the human gut microbiota and the recreated microbiota in R_c-H2_. **c**. Correlation of functional gene abundance between human gut microbiota and recreated microbiota in R_c-H2_.

## Notes

### Competing Interest Statement

The authors have declared no competing interest.

