## Supplementary material for "X-Gutor: a high-fidelity *in vitro* platform for long-term gut microbiome recovery": Fig. S1-S10

**Supplemental figure**

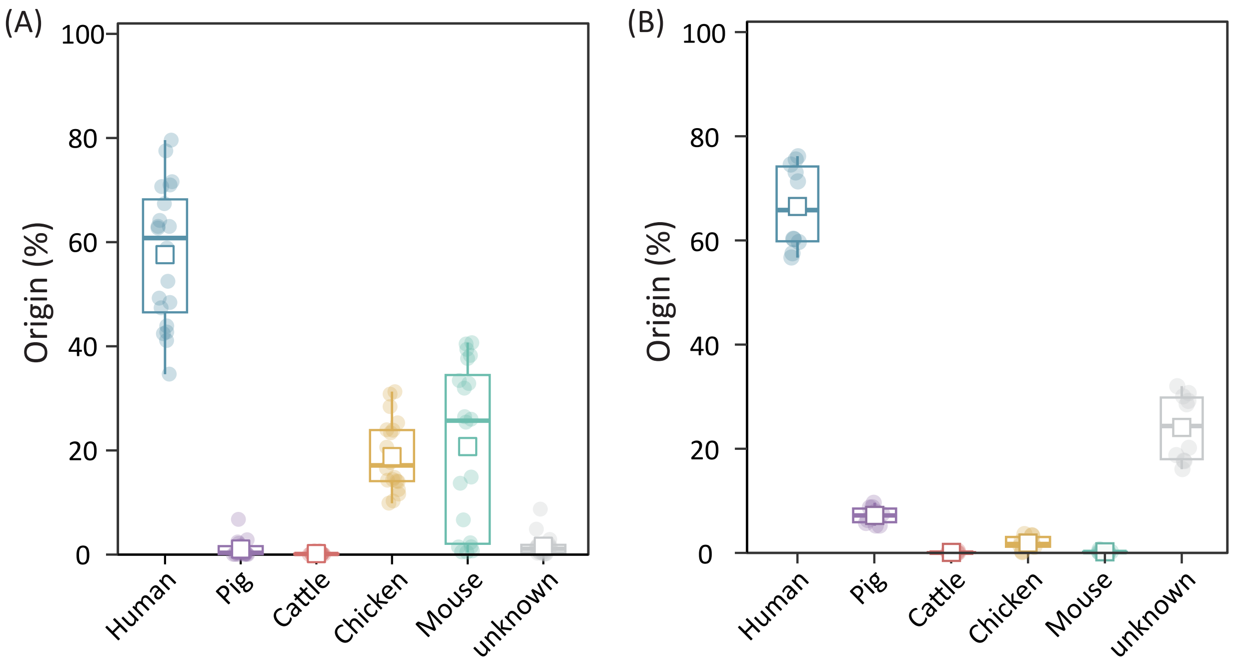


**Figure S1.** Origin estimation of recreated microbiota in the SHIME system (**a**) and gut-on-chips (**b**) based on microbial source tracking analysis.

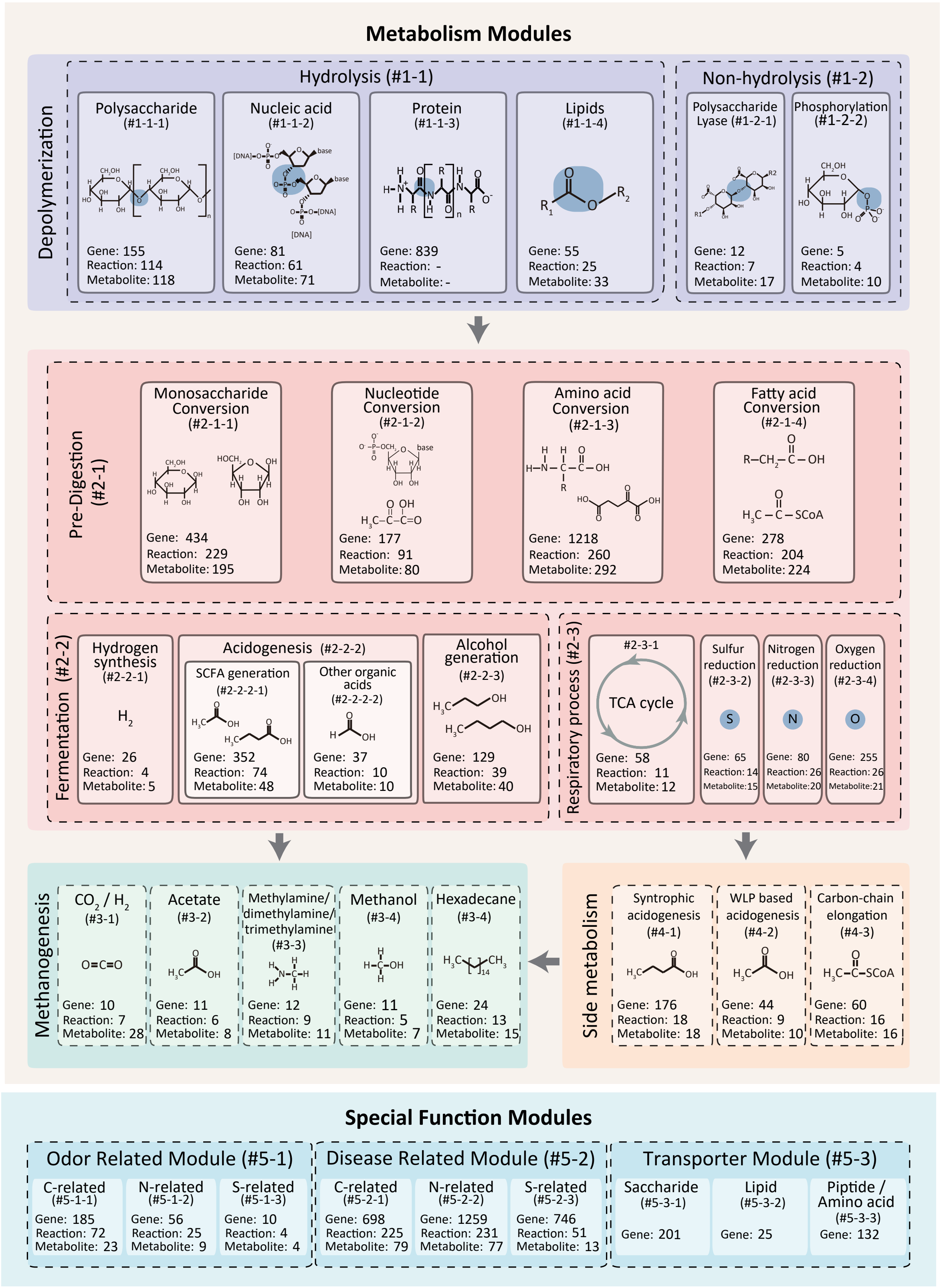


**Figure S2. General and special metabolism modules in the GutDB.** The curated metabolic pathways, genes, gene-encoded enzymes and metabolites were grouped into 5 metabolism-specific modules including organic hydrolysis, fermentation, acetogenesis, respiration, methanogenesis, and gut-microbiota-specific function modules, of which sequence data were mainly retrieved from KEGG, UniPort and MEROPS databases. The modules were numbered with pound sign (#).

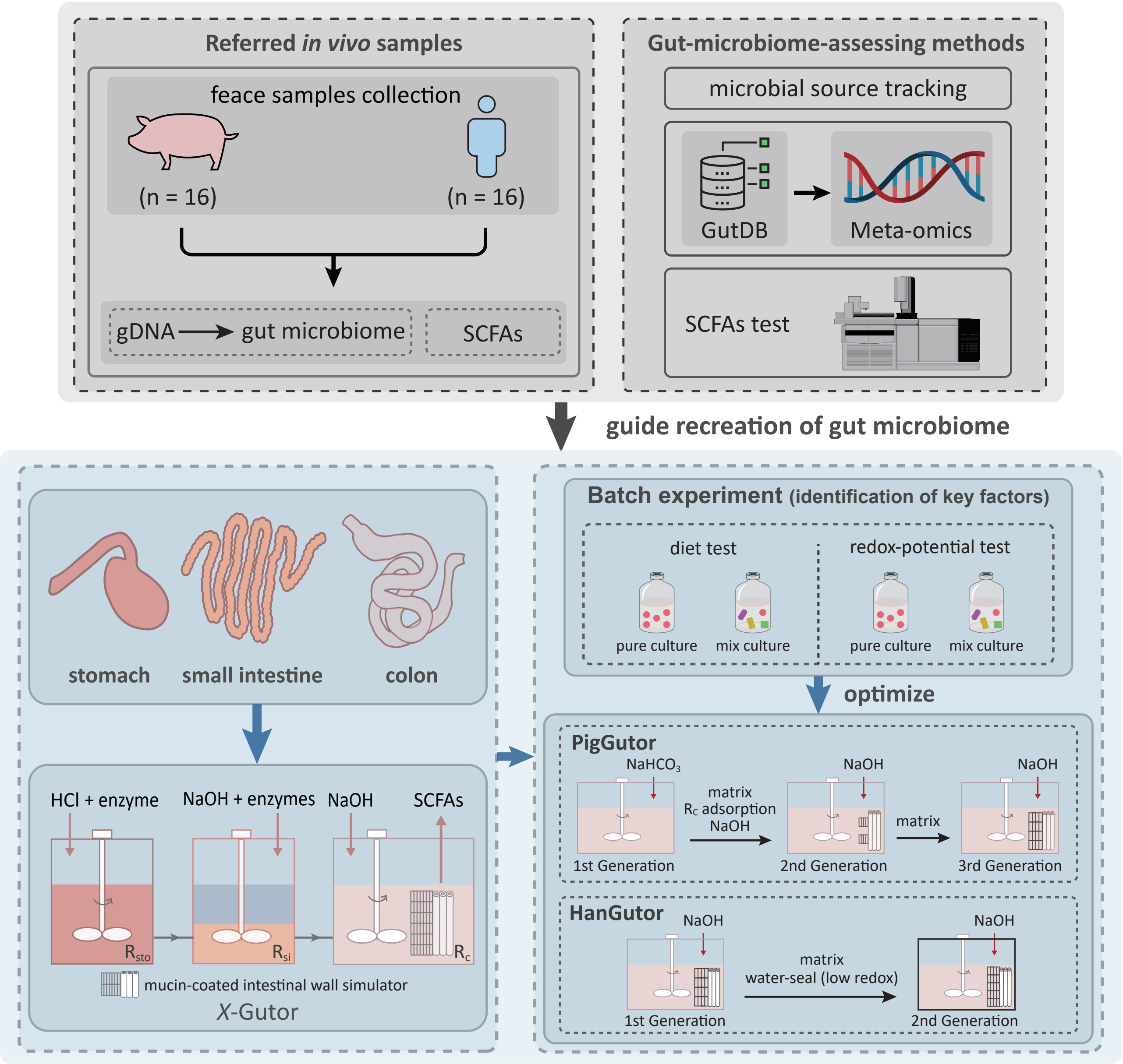


**Figure S3. Schematic diagram of the experiment.** The experiments include four parts: (1) community and SCFAs analyses of feces samples collected from swine (n = 16) and human (n = 16) donors, and the data were referred to guide the recreation of *in vitro* gut microbiome; (2) development of three methods (i.e., microbial source tracking, Gut-DB based meta-omics and SCFAs analyses) to support assessment of gut microbiome recovery at the taxonomy, gene-coded function and metabolism levels, respectively; (3) development of the *X*-Gutor to mimic monogastric digestive systems that was further extended to construction of PigGutor and HanGutor; (4) batch experiments based on both pure and mixed cultures were used to test the impact of redox potential and feed on the gut microbiome, particularly the microbial interactions.

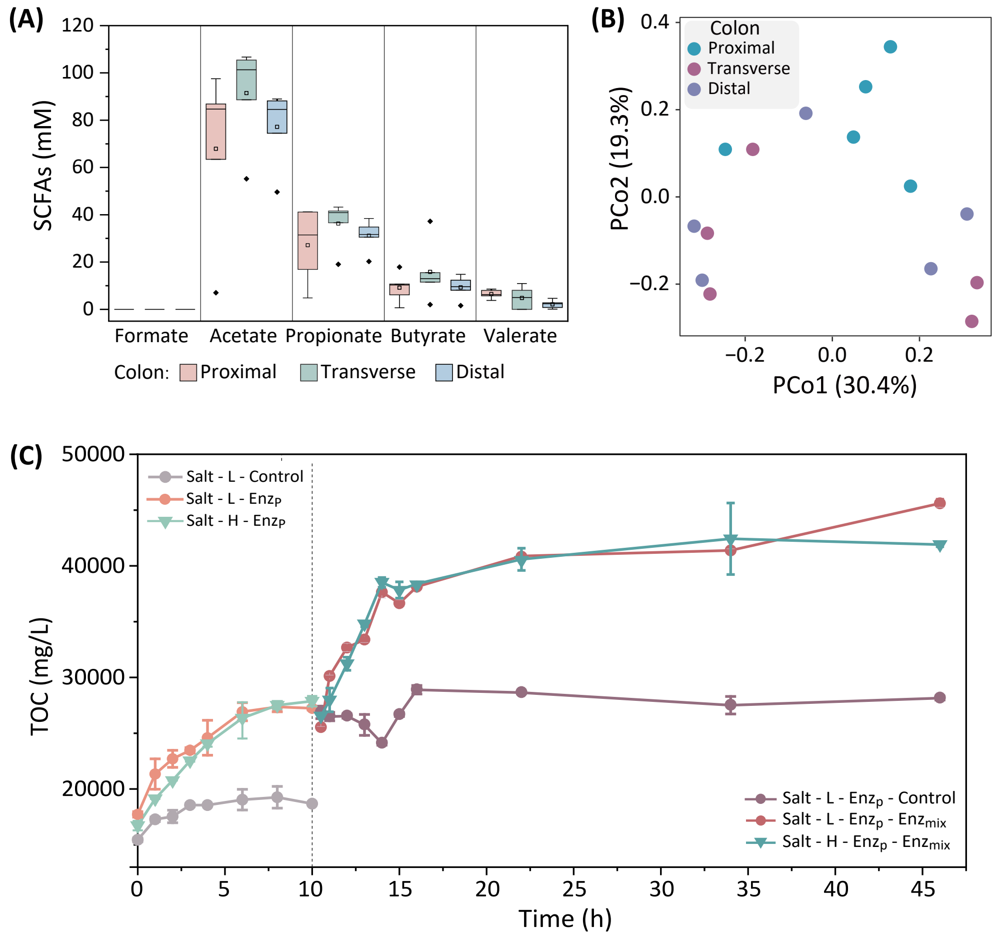


**Figure S4. Microbial and SCFAs characterization of swine gut. a**. SCFAs concentration and profile of swine colon (proximal, transverse and distal colon) samples. **b**. Principal coordinate analysis (PCoA) of the proximal, transverse and distal colon communities. **c**. Impact of high/low salinity and digestion time on digestion efficiency quantified with dissolved organic matter in stomach ($R_{sto}$) and small intestine ($R_{si}$) simulators. Feeding substrates were first digested in the Rsto for 10 hours (Stage-I, 0 – 10 hours) and then in the $R_{si}$ during subsequent 38 hours (Stage-II, 10 – 48 hours). In each stage, 3 sets of experiments were setup. In Stage-I: Salt-L-Control, control experimental set without pepsase addition and in low salt condition; Salt-L-$Ezn_P$, experimental set with pepsase addition and in low salt condition; Salt-H-$Ezn_P$, experimental set with pepsase addition and in high salt condition. In Stage-II: Salt-L-$Ezn_P$-Control, control experimental set conducted with the Salt-L-$Ezn_P$ digestion product (from Stage-I) without addition of mixed enzymes; Salt-L-EznP-$Ezn_{mix}$, experimental set conducted with the Salt-L-$Ezn_P$ digestion product with mixed enzymes amendment; Salt-H-$Ezn_P$-$Ezn_{mix}$, experimental set conducted with the Salt-H-$Ezn_P$ digestion product with mixed enzymes addition. See detailed information on salts and digestion enzymes, as well as their concentrations, in **Table S4**.

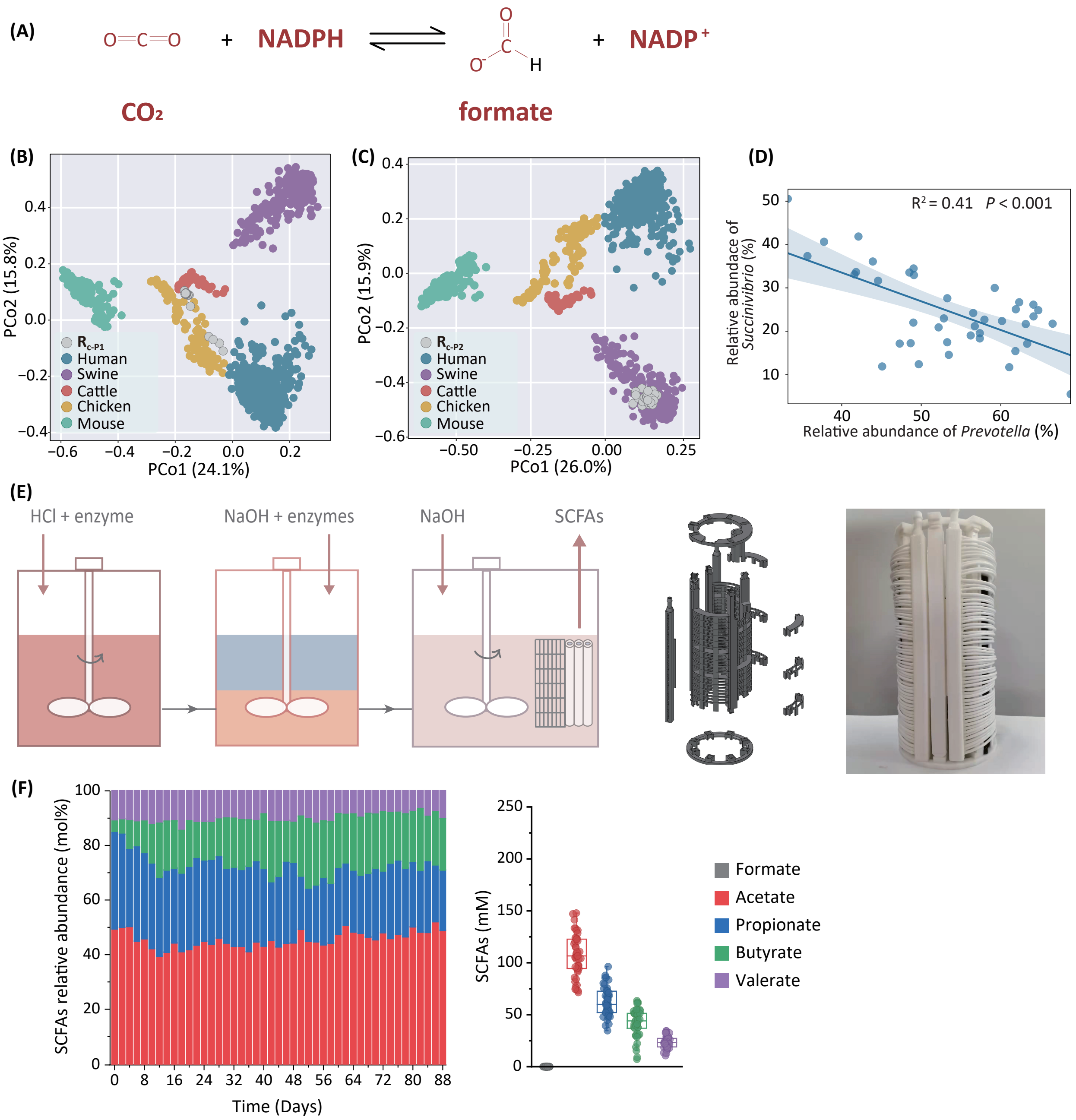


**Figure S5. Recovery improvement of *in vitro* swine gut microbiota in PigGutor. a**. Formation of formate from $CO_2$ and NADPH under mediation of NADP-dependent formate dehydrogenase. PCoA of *in vitro* swine gut communities in $R_{c\text{-}P1}$ (**b**), and $R_{c\text{-}P2}$ (**c**), together with potential source communities. **d**. Correlation between relative abundance of *Succinivibrio* and *Prevotella* (n = 40). **e**. Optimization of PigGutor by replacing $NaHCO_3$ with NaOH as a pH adjustment chemical and by adding mucin-coated intestinal wall simulator. **f**. Temporal change in SCFAs concentration and composition in the 3rd generation swine colon simulator ($R_{c\text{-}P3}$).

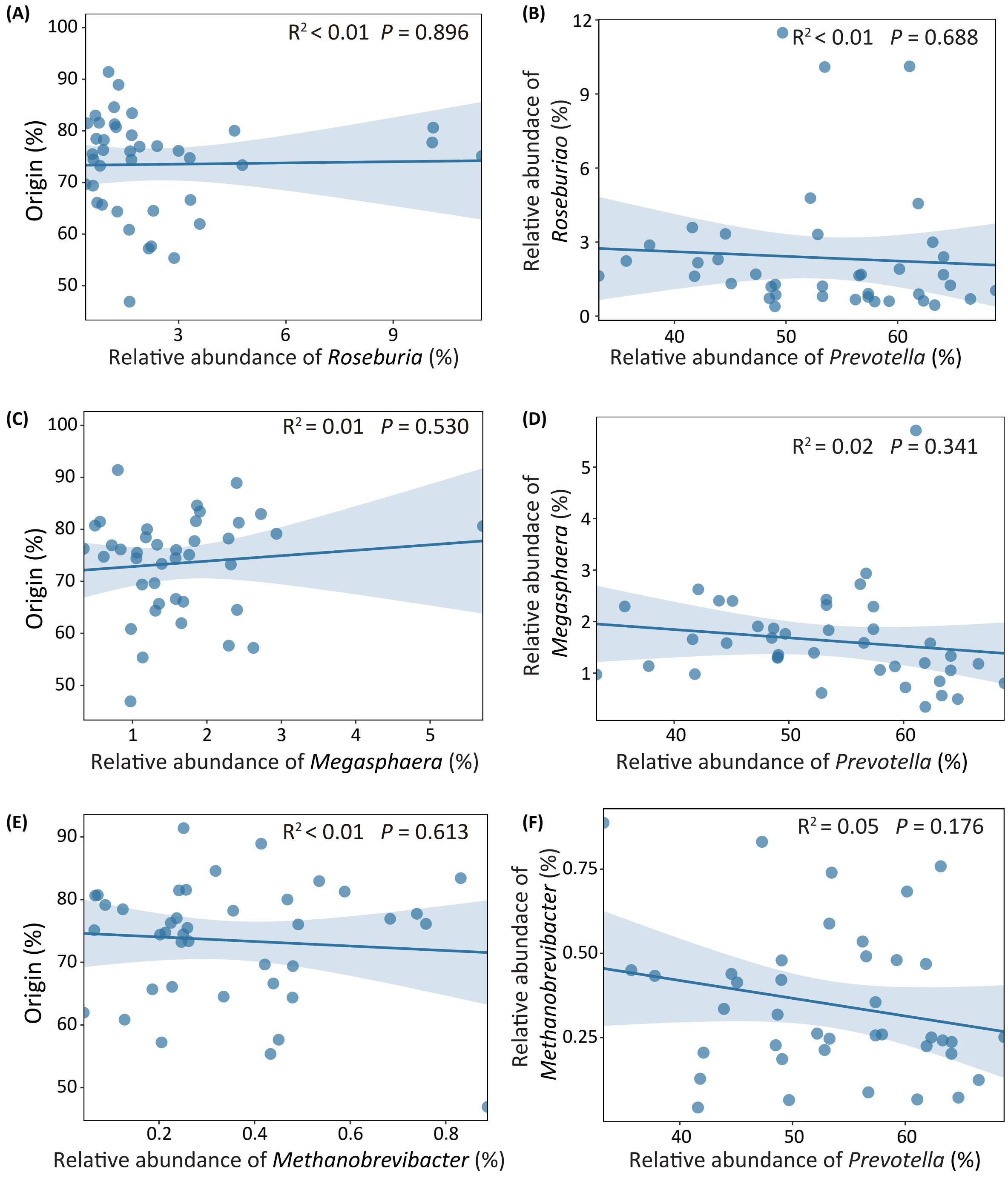


**Figure S6. Correlations of major populations' relative abundance with source-tracking-based recovery ratio of swine gut microbiota in $R_{c\text{-}P2}$ and with the relative abundance of *Prevotella*.** Correlations between the relative abundance of *Roseburia* (**a**), *Megasphaera* (**c**), and *Methanobrevibacter* (**e**) and the recovery ratio of swine gut microbiota in $R_{c\text{-}P2}$. $R^2$, coefficient of determination (n = 40). Correlations between the relative abundance of *Roseburia* (**b**), *Megasphaera* (**d**), and *Methanobrevibacter* (**f**) with the relative abundance of *Prevotella* (n = 40).

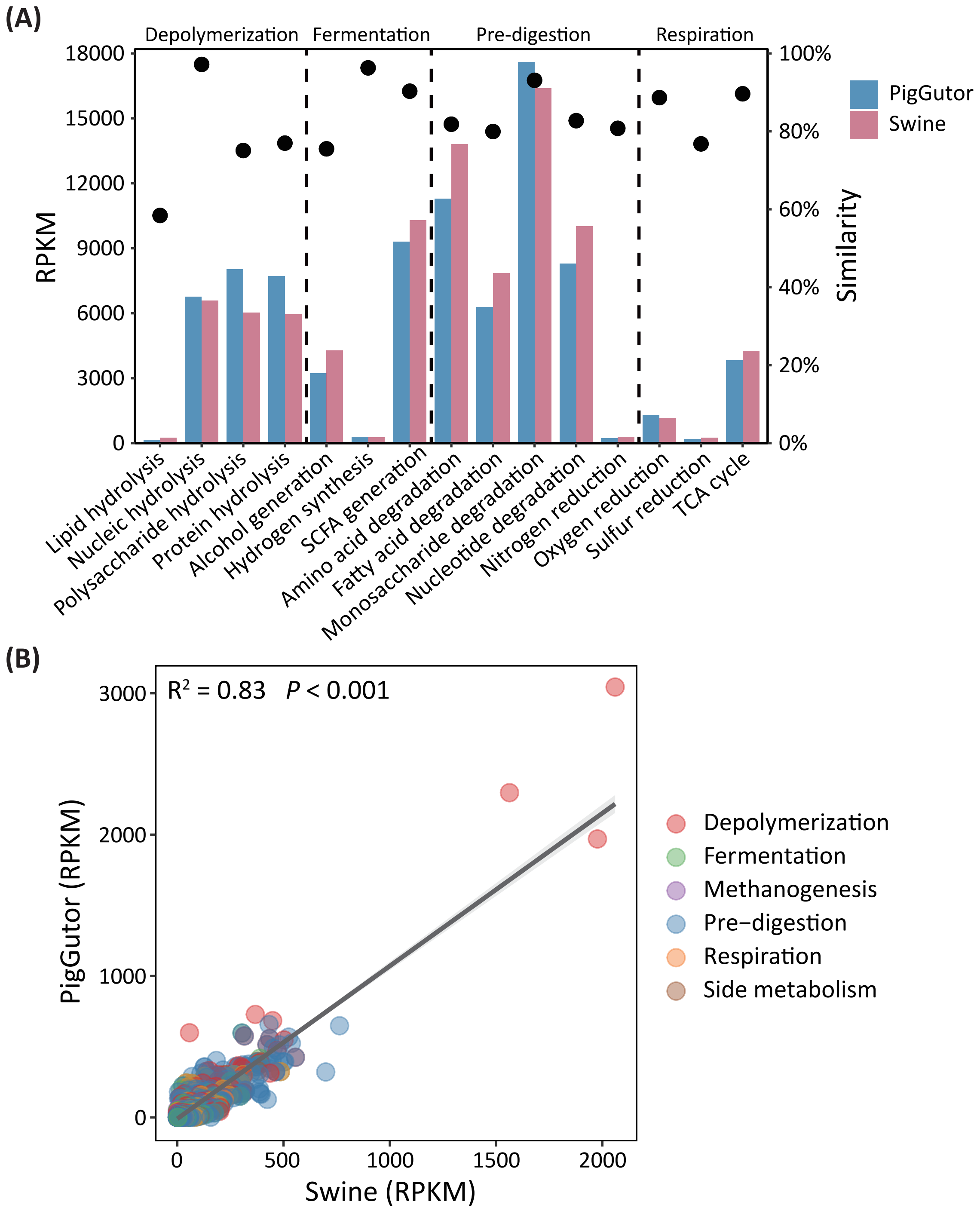


**Figure S7. Similarity comparison of metabolism modules between *in vitro* and *in vivo* swine gut microbiota based on the GutDB-supporting metagenomic analyses. a**. Similarity of major metabolism modules between the swine gut microbiota and the recreated microbiota in $R_{c\text{-}P3}$. **b**. Correlation of functional gene abundance between swine gut microbiota and recreated microbiota in $R_{c\text{-}P3}$.

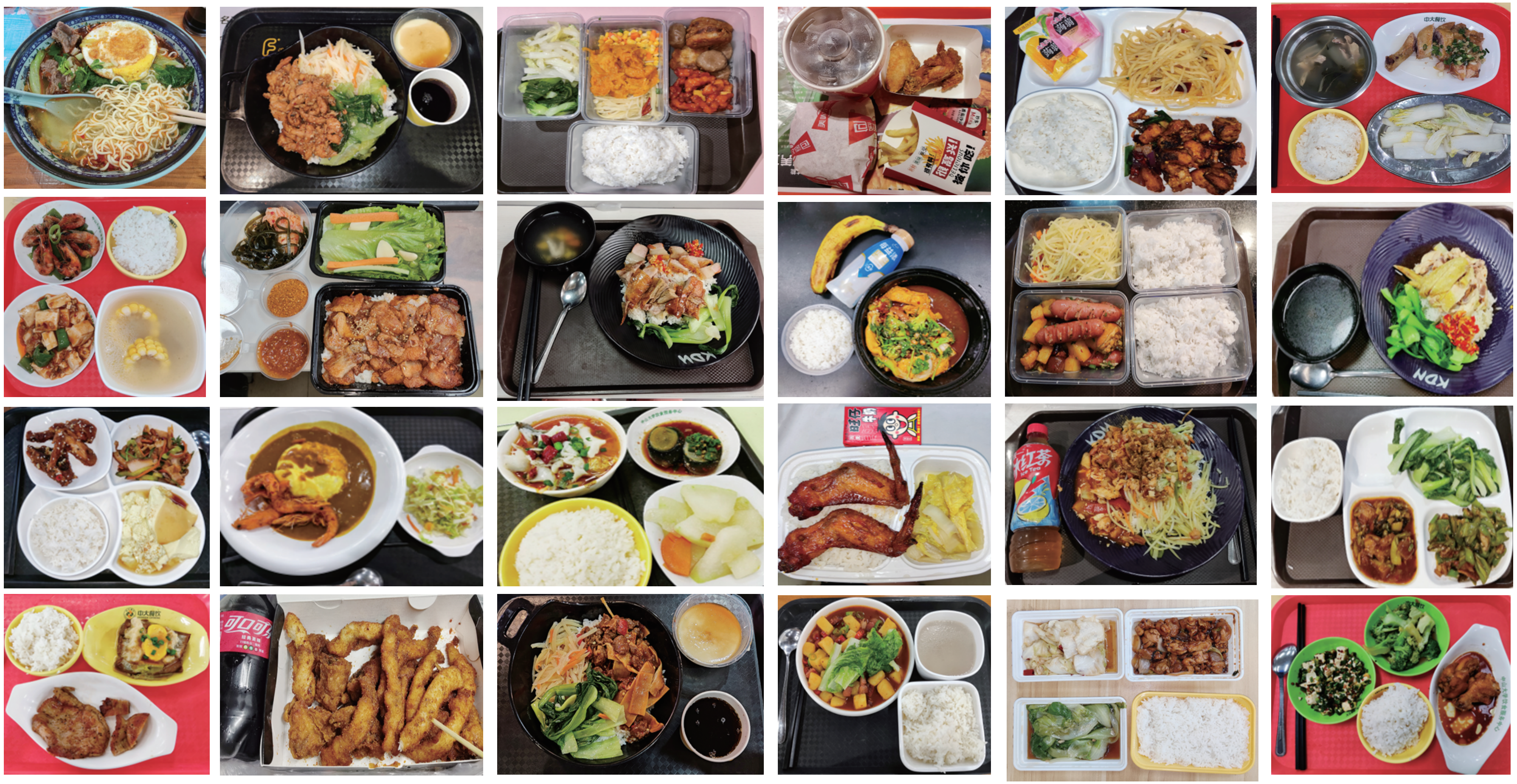

**Figure S8. Foods used as feedstock for the HanGutor.**

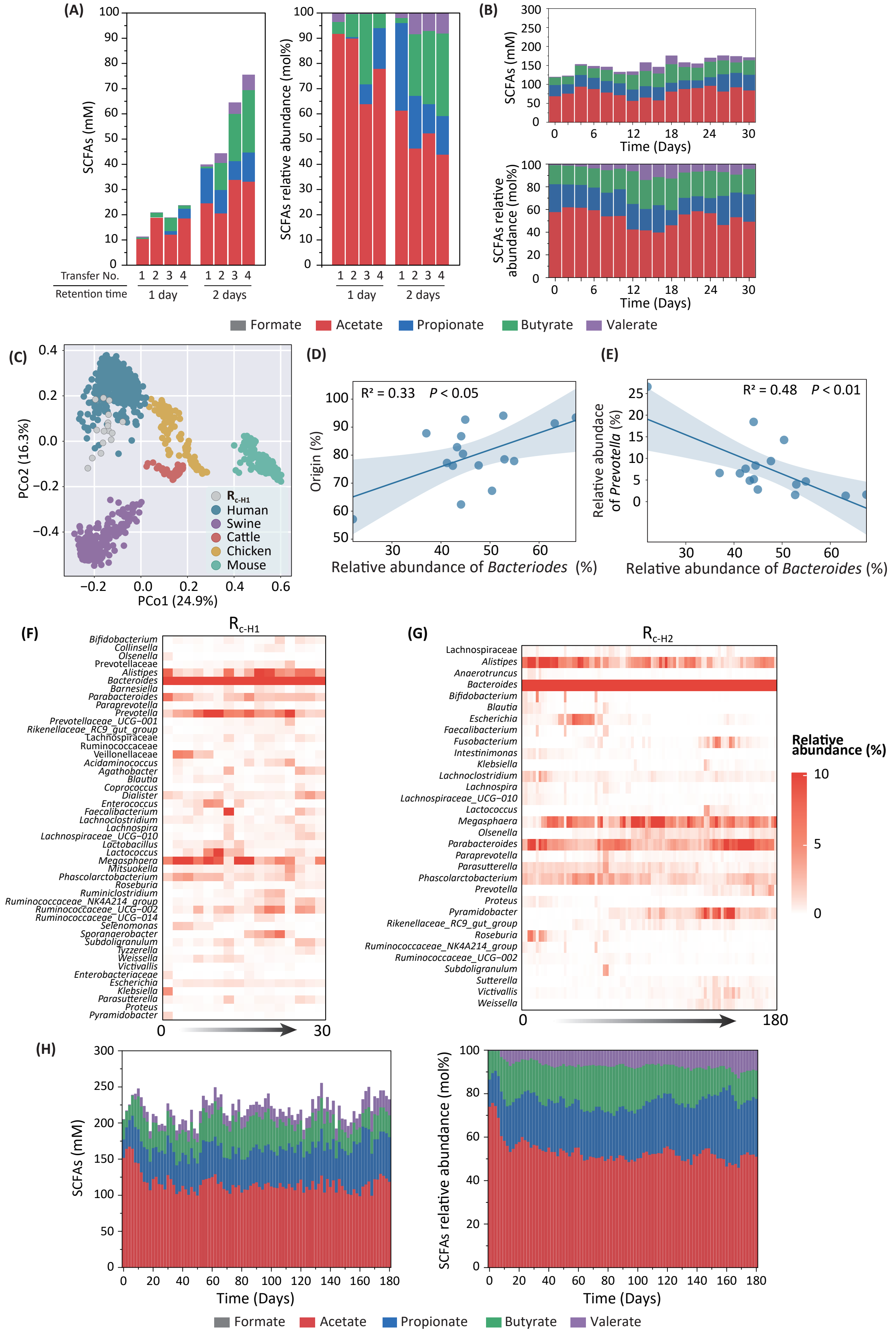


**Figure S9. Characterization of recreated human gut microbiota in HanGutor. a**. Impact of digestate retention time on SCFAs generation. **b**. Temporal changes of SCFAs concentration and composition in $R_{c\text{-}H1}$. **c**. PCoA of *in vitro* human gut communities in $R_{c\text{-}H1}$, together with potential source communities. **d**. Correlation between the relative abundance of *Bacteroides* and the microbial-source-tracking-calculated recovery ratio of recreated microbiota in $R_{c\text{-}H1}$ (n=16). **e**. Correlation between the relative abundance of *Bacteroides* and *Prevotella* in $R_{c\text{-}H1}$ (n=16). **f**. Temporal change in major populations (average relative abundance > 0.1%) at the genus level in $R_{c\text{-}H1}$. **g**. Temporal change in major populations (average relative abundance > 0.1%) at the genus level in $R_{c\text{-}H2}$. **h**. Temporal changes of SCFAs concentration and composition in $R_{c\text{-}H2}$.

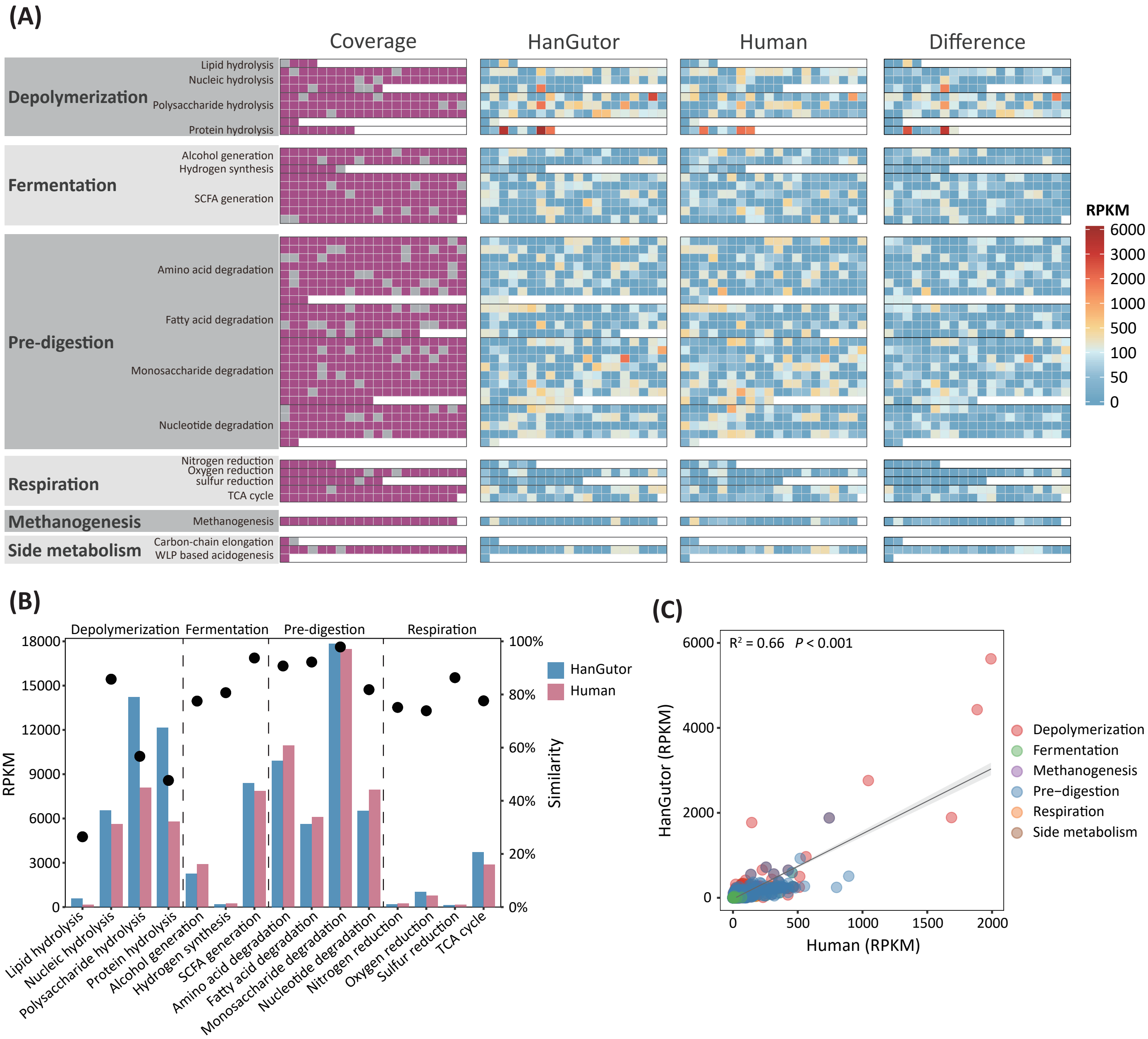


**Figure S10. Comparison of *in vitro* and *in vivo* human gut microbiota at the functional level. a**. Coverage and functional profile comparison of the recreated microbiota in $R_{c\text{-}H2}$ and *in vivo* human gut microbiota based on GutDB-supported metagenomic analysis. **b**. Similarity of major metabolism modules between the human gut microbiota and the recreated microbiota in $R_{c\text{-}H2}$. **c**. Correlation of functional gene abundance between human gut microbiota and recreated microbiota in $R_{c\text{-}H2}$.
