## Supplementary material for "X-Gutor: a high-fidelity *in vitro* platform for long-term gut microbiome recovery": Table S1-S6

### Supplemental Tables

**Table S1.** Accession numbers of previously sequencing data for microbial source tracking analysis.

| Source types | Accession number | Samples | Hypervariable region | Platform | Reference |
|---|---|---|---|---|---|
| Cattle | - | 93 | V4-V5 | Illumina HiSeq 2500 | This study |
| Chicken | PRJEB32628 | 21 | V3-V4 | Illumina Miseq | Kempf et al.[S1] |
| | PRJNA606870 | 6 | V3-V4 | Illumina HiSeq 2500 | Wu et al. [S2] |
| | PRJNA552082 | 69 | V3-V4 | Illumina HiSeq 2000 | Robinson et al. [S3] |
| | PRJNA633979 | 37 | V3-V4 | Illumina Miseq | Tolnai et al. [S4] |
| Human | PRJNA542910 | 117 | V4 | Illumina Miseq | Grembi et a.. [S5] |
| | PRJNA605207 | 89 | V3-V4 | Illumina Miseq | Serrano et al. [S6] |
| | PRJNA480547 | 318 | V3-V4 | Illumina Miseq | Wan et al. [S7] |
| Mouse | PRJNA836043 | 31 | V4 | Illumina Miseq | Zhang et al. [S8] |
| | PRJNA682275 | 118 | V4 | Illumina MiSeq | Diamond et al. [S9] |
| Swine | PRJNA531671 | 55 | V4 | Illumina MiSeq | Wang et al. [S10] |
| | PRJNA356465 | 215 | V4 | Illumina MiSeq | Yang et al. [S11] |

**Table S2.** Basic information of human feces donors.

| Number | Gender | Age (year old) | Height (m) | Weight (kg) | BMI | Weight status | Defecation interval (d) | Moisture content (%) |
|---|---|---|---|---|---|---|---|---|
| H-1[§] | F | 23 | 1.63 | 41 | 15.7 | underweight | 3 | 71 |
| H-2 | M | 32 | 1.76 | 71.5 | 23.1 | healthy | 1 | 91 |
| H-3 | F | 25 | 1.62 | 52.5 | 20 | healthy | 1 | 85 |
| H-4 | M | 24 | 1.83 | 102.5 | 30.6 | obese | 1 | 79 |
| H-5 | M | 22 | 1.74 | 68 | 22.5 | healthy | 1 | 76 |
| H-6 | F | 12 | 1.58 | 42 | 16.8 | underweight | 5 | 73 |
| H-7 | F | 12 | 1.55 | 40 | 16.6 | underweight | 4 | 64 |
| H-8 | F | 22 | 1.55 | 44 | 18.3 | underweight | 1 | 80 |
| H-9 | M | 30 | 1.65 | 58 | 21.3 | healthy | 1 | 84 |
| H-10 | F | 28 | 1.58 | 47 | 18.8 | healthy | 2.5 | 78 |
| H-11 | F | 26 | 1.57 | 50 | 22.3 | healthy | 2 | 81 |
| H-12 | F | 26 | 1.57 | 43 | 17.4 | underweight | 1.5 | 89 |
| H-13 | F | 24 | 1.6 | 47 | 18.4 | healthy | 1.5 | 75 |
| H-14 | M | 50 | 1.73 | 51 | 17 | underweight | 3 | 72 |
| H-15 | M | 50 | 1.73 | 52.5 | 17.5 | underweight | 2.5 | 69 |
| H-16 | F | 28 | 1.6 | 52.5 | 20.5 | healthy | 2 | 59 |

§ feces donor for setup of the HanGutor

**Table S3.** Basic information of swine digestate/feces donors.

| Sample type | Segment | Number of samples | Total sample number | Raising time (month) | Weight (kg) |
|---|---|---|---|---|---|
| swine feces | / | 16 | 16 | 6 | 90-110 |
| swine digestate | proximal, transverse, and distal colon | 5 | 15 | 6 | 90-110 |

**Table S4.** Parameters for setup and operation of the *in vitro* gut microbiome culturing models.

| | | | PigGutor | | | HanGutor | |
|---|---|---|---|---|---|---|---|
| | | | G1 | G2 | G3 | G1 | G2 |
| | | Feed | corn : soybean : chicken-powder : wheat-bran : premix (50:20:20:5:5; w/w) | | | meat : vegetable : starches : fruit (22:38:25:15; w/w) | |
| Pretreatment | $R_{sto}$ | pH | 2 | | | 2 | |
| | | enzyme | pepsase (738 U/mL) | | | pepsase (738 U/mL) | |
| | | salinity | low salinity | | | low salinity | |
| | | retention time (h) | 10 | | | 10 | |
| | $R_{si}$ | pH | 7 | | | 7 | |
| | | enzyme | amylase (221 U/mL), lipase (3 U/mL), chymotrypsin (9 U/mL), trypsin (69 U/mL) | | | amylase (221 U/mL), lipase (3 U/mL), chymotrypsin (9 U/mL), trypsin (69 U/mL) | |
| | | salinity† | low salinity (0 mol/L NaCl) | | | low salinity (0 mol/L NaCl) | |
| | | retention time (h) | 12 | | | 12 | |
| Digestion | $R_c$ | identity | $R_{c-P1}$ | $R_{c-P2}$ | $R_{c-P3}$ | $R_{c-H1}$ | $R_{c-H2}$ |
| | | pH control | 6.1-6.4 $NaHCO_3$ | 6.1-6.4 NaOH | 6.1-6.4 NaOH | 6.4-6.8 NaOH | 6.4-6.8 NaOH |
| | | water seal | no (anoxic) | | | no (anoxic) | yes (obligate anaerobic) |
| | | retention time (days) | 1.25 | | | 2 | |
| | | assimilation | no | yes | yes | yes | yes |
| | | biofilm matrix (‰, v/v) | no | 5 | 20 | 20 | 30 |

§ pH was adjusted to 5.0 via the pH controller

† high salinity (0.4 mol/L NaCl); low salinity (0 mol/L NaCl)

**Table S5.** ANOVA test on the functional gene abundances in metabolism modules between PigGutor and swine feces and between HanGutor and human feces.

| Module | *P* value | |
|---|---|---|
| | PigGutor VS. Swine | HanGutor VS. Human |
| Depolymerization | 0.497 | 0.121 |
| Fermentation | 0.329 | 0.924 |
| Pre-digestion | 0.307 | 0.567 |
| Respiration | 0.742 | 0.350 |
| Methanogenesis | 0.244 | 0.654 |
| Side metabolism | 0.324 | 0.353 |

**Table S6.** Mice diet compositions.

| Diet formulas | Diverse diet | | Monotonous diet | |
|---|---|---|---|---|
| Product | gm% | kcal% | gm% | kcal% |
| Protein | 20.3 | 20.5 | 0.3 | 0.3 |
| Carbohydrate | 51.7 | 52.2 | 95.0 | 99.7 |
| Fat | 12 | 27.3 | 0.0 | 0.0 |
| Total | | 100.0 | | 100.0 |
| Kcal/gm | 4.00 | | 3.85 | |
| Ingredient | gm | Kcal | gm | Kcal |
| Corn Starch | 335 | 1340 | 767.5 | 3070 |
| Casein | 200 | 800 | 0 | 0 |
| Maltodextrin | 102 | 408 | 102 | 408 |
| Sucrose | 80 | 320 | 80 | 320 |
| Soybean Oil | 120 | 1080 | 0 | 0 |
| Cellulose | 112.5 | 0 | 0 | 0 |
| Mineral Mix S10022G | 35 | 0 | 35 | 0 |
| Vitamin Mix V10037 | 10 | 40 | 10 | 40 |
| L-Cystine | 3 | 12 | 3 | 12 |
| Choline Bitartrate | 2.5 | 0 | 2.5 | 0 |

### Refferences

[S1] Kempf, F. *et al*. Gut microbiota composition before infection determines the Salmonella super-and low-shedder phenotypes in chicken. *Microb. Biotechnol.* **13**, 1611-1630 ( 2020).

[S2] Wu, S., Chen, X., Li, T., Ren, H., Zheng, L. and Yang, X. Changes in the gut microbiota mediate the differential regulatory effects of two glucose oxidases produced by Aspergillus niger and Penicillium amagasakiense on the meat quality and growth performance of broilers. *J. Anim. Sci. Biotechnol*. **11**, 1-13 (2020).

[S3] Robinson, K. *et al*. Differential impact of subtherapeutic antibiotics and ionophores on intestinal microbiota of broilers. *Microorganisms*, **7**, 282 (2019).

[S4] Tolnai, E. *et al*. Nutraceuticals induced changes in the broiler gastrointestinal tract microbiota. *MSystems*, **6**, 10-1128 (2021).

[S5] Grembi, J.A., Nguyen, L.H., Haggerty, T.D., Gardner, C.D., Holmes, S.P. and Parsonnet, J. Gut microbiota plasticity is correlated with sustained weight loss on a low-carb or low-fat dietary intervention. *Sci. Rep.* **10**, 1405 (2020).

[S6] Serrano, J. *et al*. High-dose saccharin supplementation does not induce gut microbiota changes or glucose intolerance in healthy humans and mice. *Microbiome*, **9**, 1-18 (2021).

[S7] Wan, Y. *et al*. Effects of dietary fat on gut microbiota and faecal metabolites, and their relationship with cardiometabolic risk factors: a 6-month randomised controlled-feeding trial. *Gut*, **68**, 1417-1429 (2019).

[S8] Zhang, Z. *et al*. Gracilariopsis lemaneiformis Polysaccharide Attenuates D-Galactose–Induced Aging of Mice by Regulating Oxidative Stress and Gut Microbiota. *Front. Mar. Sci.* **9**, 932147 (2022).

[S9] Diamond, E., Hewlett, K., Penumutchu, S., Belenky, A. and Belenky, P. Coffee consumption modulates amoxicillin-induced dysbiosis in the murine gut microbiome. *Front. Microbiol.* **12**, 637282 (2021).

[S10] Wang, X. *et al*. Longitudinal investigation of the swine gut microbiome from birth to market reveals stage and growth performance associated bacteria. *Microbiome*, **7**, 1-18 (2019).

[S11] Yang, H. *et al*. Evaluating the profound effect of gut microbiome on host appetite in pigs. *BMC Microbiol.* **18**, 1-10 (2018).
